# Representation of locomotive action affordances in human behavior, brains and deep neural networks

**DOI:** 10.1101/2024.05.15.594298

**Authors:** Clemens G. Bartnik, Christina Sartzetaki, Abel Puigseslloses Sanchez, Elijah Molenkamp, Steven Bommer, Nikolina Vukšić, Iris I. A. Groen

## Abstract

To decide how to move around the world, we must determine which locomotive actions (e.g., walking, swimming, or climbing) are afforded by the immediate visual environment. The neural basis of our ability to recognize locomotive affordances is unknown. Here, we compare human behavioral annotations, functional magnetic resonance imaging (fMRI) measurements, and deep neural network (DNN) activations to both indoor and outdoor real-world images to demonstrate that human visual cortex represents locomotive action affordances in complex visual scenes. Hierarchical clustering of behavioral annotations of six possible locomotive actions show that humans group environments into distinct affordance clusters using at least three separate dimensions. Representational similarity analysis of multi-voxel fMRI responses in scene-selective visual cortex shows that perceived locomotive affordances are represented independently from other scene properties such as objects, surface materials, scene category or global properties, and independent of the task performed in the scanner. Visual feature activations from DNNs trained on object or scene classification as well as a range of other visual understanding tasks correlate comparatively lower with behavioral and neural representations of locomotive affordances than with object representations. Training DNNs directly on affordance labels or using affordance-centered language embeddings increases alignment with human behavior, but none of the tested models fully captures locomotive action affordance perception. These results uncover a new type of representation in the human brain that reflects locomotive action affordances.

**Significance:** To navigate the world around us, we can use different actions, such as walking, swimming or climbing. How does our brain compute and represent such locomotive action affordances? Here, we show that activation patterns in high-level visual regions in the human brain represent information about affordances independent of other visual elements such as surface materials and objects, and do so in an automatic manner. We also demonstrate that commonly used models of visual processing in human brains, namely object- and scene-classification trained deep neural networks, do not strongly represent this information. Our results suggest that locomotive action affordance perception in scenes relies on specialized neural representations different from those used for other visual understanding tasks.

## Introduction

Humans navigate their local environments with remarkable ease, whether cycling through busy city streets or hiking along rugged trails. This highlights the ability of the human brain to not only effortlessly process visual information, but also identify appropriate locomotive actions from a wide range of potential actions. The recognition of ecologically relevant behaviors in the immediate environment is commonly referred to as *affordance perception* (Gibson, 1977). Theoretical frameworks of affordance perception are long-standing in fields such as ecological psychology (Gibson, 1977; Rietveld and Kiverstein, 2014; Osiurak et al., 2017), but the neural mechanisms mediating this ability during scene perception are only just beginning to be explored.

While initial research on scene perception emphasized scene-defining objects and their relationships (e.g. Biederman et al., 1982; Aminoff et al., 2013), later work included global features such as spatial layout (Oliva and Torralba, 2001), as well as affordance-related concepts such as navigability (Greene and Oliva, 2009) (see Bartnik and Groen, 2023, for a review). Functional magnetic resonance imaging (fMRI) has identified three brain regions involved in processing complex visual scenes (Epstein and Baker, 2019) — the Parahippocampal Place Area (PPA), Occipital Place Area (OPA), and Medial Place Area (MPA, also known as retrosplenial cortex or RSC; Silson et al., 2016). Of these, PPA has been consistently associated with the representation of global spatial layout of scenes (Epstein and Kanwisher, 1998; Kravitz et al., 2011; Park et al., 2011, 2015), while a pioneering study by Bonner and Epstein (2017) showed that OPA represents navigable space, in the form of pathways that afford walking. Subsequent analyses showed that these pathway representations in OPA aligned with representations of extended surfaces and floor elements in convolutional neural networks (CNNs) trained on scene recognition or scene segmentation (Bonner and Epstein, 2018; Dwivedi et al., 2021b). These and other findings (Park and Park, 2020; Persichetti and Dilks, 2016; Dilks et al., 2011; Julian et al., 2016; Kamps et al., 2016b,a) gave rise to the hypothesis that OPA plays a prominent role in the computation of navigational scene affordances in particular.

Further empirical evidence of an important role of affordances in scene perception was found by Greene et al. (2016), who showed that diverse actions, such as playing sports and transportation, strongly affect human scene categorization of a wide range of environments in a large-scale scene dataset. Groen et al. (2018) replicated these behavioral findings, but found no evidence for representation of broadly defined affordances in scene-selective cortex, nor in object- or scene- classification trained CNNs. These results thus argue against a role of scene-selective brain regions, including OPA, in representing affordances. The discrepancy in findings with e.g. Bonner and Epstein (2017) may be due to differences in the types of stimuli used (diverse environments versus indoor scenes) and to how affordances were operationalized (potential action labels versus navigable pathways). One intriguing hypothesis that could potentially unify these findings is that scene-selective regions represent only a distinct subset of action affordances, namely those that pertain specifically to navigation. Here, we test if scene-selective regions differentiate visual environments based on whether they afford locomotive actions *other* than walking, such as cycling, driving or swimming.

Given OPA’s hypothesized role in navigational affordance perception, we might expect such locomotive action affordances to be primarily encoded in OPA; however, other work linking spatial scene and object properties with human interaction possibilities (Bainbridge and Oliva, 2015; Josephs and Konkle, 2020) also implicates other visual brain regions. *A priori*, it seems likely that disambiguating different locomotive action affordances requires perceiving not only spatial properties of scenes (e.g., the presence of navigable surfaces such floors or roads) but also surface materials (asphalt for driving or cycling) and contained objects (rocks for climbing, a body of water for swimming), i.e., a variety of diagnostic features (Groen et al., 2017). Such properties may be encoded not only in OPA but also PPA, which represents not only the spatial layout of scenes, but also textures (Henriksson et al., 2019; Cant and Xu, 2012) and objects (Janzen and van Turennout, 2004; Marchette et al., 2015; Harel et al., 2013). However, it is unclear which, if any, combination of these properties is sufficient to predict the perceived affordances of a scene, and their corresponding representation in the human brain, or whether additional (possibly non-visual) information is involved. Finally, neural representation of action-related information has been shown to be sensitive to task context (e.g., Bracci et al., 2017), but prior studies did not probe participants to actively report perceived scene affordances during fMRI measurements, which may impact neural representations of diagnostic features.

Here, we investigated if and how humans represent locomotive action affordances by testing how well various scene properties predict action affordance representation as assessed with both behavioral and fMRI measurements. Using a novel set of natural scenes spanning six common locomotive affordances, we test the hypothesis that human scene-selective cortex represents action possibilities that pertain to locomotion. We furthermore assert whether locomotive affordance representations are task-dependent by comparing neural activation patterns during tasks that require participants to explicitly report locomotive affordances, versus tasks that do not. To probe the neural computations underlying affordance representations, we examine how well these representations can be predicted by features extracted from a wide variety of deep neural network models (DNNs), including image, video and multi-modal vision-language models, and we explore how well linguistic descriptions from large language models (LLMs) can capture these representations. Collectively, our results provide novel evidence for locomotive action affordance representation in scene-selective cortex that is not fully captured by other scene properties, DNNs, or LLMs.

## Results

To investigate whether the human brain represents locomotive action possibilities in real-world scenes, we collected and curated a novel set of scene images, spanning indoor, outdoor man-made, and outdoor natural environments (**Fig. 1A**). Human participants (*n* = 152) then annotated these images on six different actions that people can use to move in the immediate environment (**Fig. 1B**, left). The images were also annotated on four other types of visual scene properties thought to be important for scene perception (materials, scene category, objects, and global properties; **Fig. 1B**, right). We then applied representational similarity analysis (RSA, Kriegeskorte (2008)) by computing, for each pair of images, the dissimilarity in the annotations of locomotive action affordances as well as annotations of the four different visual scene properties, resulting in five representational dissimilarity matrices (RDMs), that we then compared with one another. In RSA, high dissimilarity between a pair of images in the RDM indicates these images differ strongly on the annotated property (e.g., one affording swimming while the other affords walking), while low dissimilarity indicates the images are perceived as having similar properties (e.g., both afford swimming). Furthermore, a high correlation between RDMs of two different scene properties indicates that these properties overlap, while a low correlation is indicative of independent representations of each property. We also used RSA on a subset of these images and tasks in an fMRI experiment, to test for locomotive action affordance representations in neural responses in scene-selective cortex, and on visual feature activations extracted from a variety of DNNs (**Fig. 1C**). Here, we interpret a significant RSA correlation between fMRI response patterns with the human behavioral annotations for a given scene property as evidence for neural representation of that property, while a high RSA correlation between DNN feature activations and human behavior or brain responses indicates that the DNN adequately captures the human representations, i.e. is considered to be ”representationally aligned” (Sucholutsky et al., 2023).

**Figure 1:**
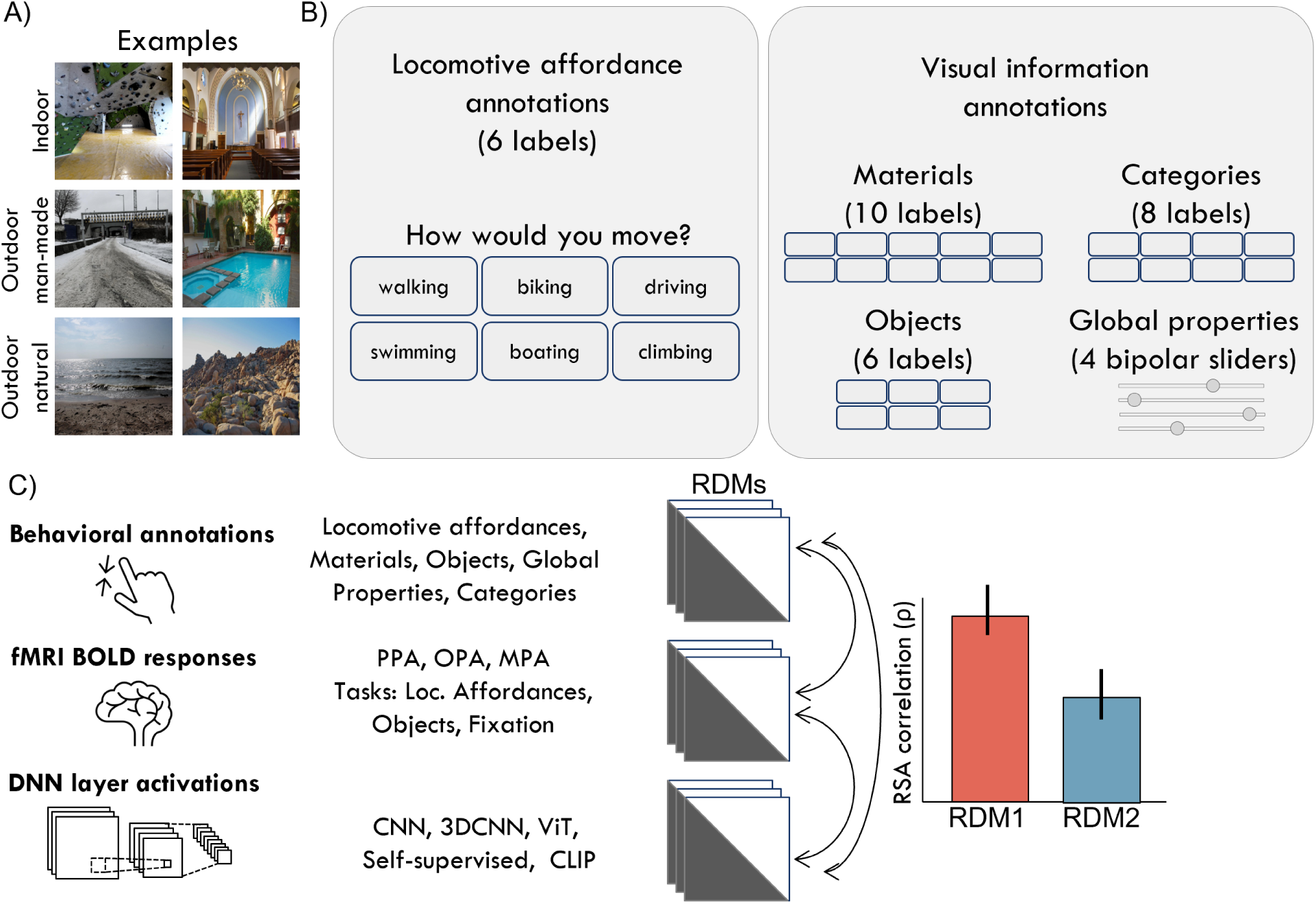
Stimuli, behavioral tasks and experimental design. **(A)** Examples from our newly curated stimulus set to measure locomotive action affordance representation consisting of 231 high-resolution photographs (collected from Flickr containing (CC0 1.0, CC BY 2.0 and CC BY-SA 4.0 licensed), evenly distributed across indoor, outdoor man-made, and outdoor natural environments. **(B)** Overview of the five labeling tasks used to collect behavioral annotations. In addition to annotating six locomotive action affordances depicted on the left, participants also annotated the scenes on perceived materials (labels: Water, Wood, Vegetation, Dirt/Soil, Sand, Stone/Concrete, Snow/Ice, Pavement, Carpet, Metal), scene category (labels: Room, Inside city, Hallway, Forest, Desert, Open country, Mountain, Coast/River), objects (labels: Building/Wall, Tree/Plant, Road/Street, Furniture, Body of water, Rocks/Stones) and global properties (Man-made/Natural, Open/Closed, Near/Far, Navigable/Non-navigable). **(C)** In addition to behavioral annotations, multi-voxel activity patterns were measured in three scene-selective brain regions of interest (ROIs): PPA, OPA and MPA, under three different task instructions (locomotive action affordance labeling; object labeling; orthogonal task at fixation). We also extracted layer activations to this new stimulus set from a variety of DNNs trained with different task objectives. Representational similarity analysis (RSA) was then used to compare the resulting representational spaces, by computing representational dissimilarity matrices (RDMs) based on pairwise differences in responses between images for each type of measurement, and correlating those to one another.

### Locomotive action affordances form a distinct representational space

To understand how humans represent locomotive action affordances, we sorted the RDM obtained from the affordance annotation task in two ways: by the type of environment (indoor, outdoor-manmade, outdoor-natural; **Fig. 2A**, left) and after applying hierarchical clustering (see Materials and Methods; **Fig. 2A**, right). The RDM sorted on environment type shows that indoor scenes are most similar to one another in terms of the locomotive actions they afford, while outdoor scenes are more dissimilar. This is likely because most indoor environments primarily afford walking, while outdoor environments typically afford multiple actions (e.g., walking and cycling). Indeed, hierarchical clustering of the affordance RDM yields at least four major clusters, the biggest of which refers to walking, mostly containing indoor scenes (**Fig. 2A**, right). However, this cluster also contains scene images from other environments, and the remaining clusters also span multiple environment types, indicating there is no one-to-one mapping of environment type to locomotive affordance. Indeed, conceptual RDMs of environment type that explicitly separate the image set into distinct superordinate classes yield only modest correlations with the affordance RDM (indoor/outdoor-natural/outdoor-manmade: *ρ* = 0.24; indoor/ outdoor: *ρ* = 0.12; manmade/ natural: *ρ* = 0.23).

**Figure 2:**
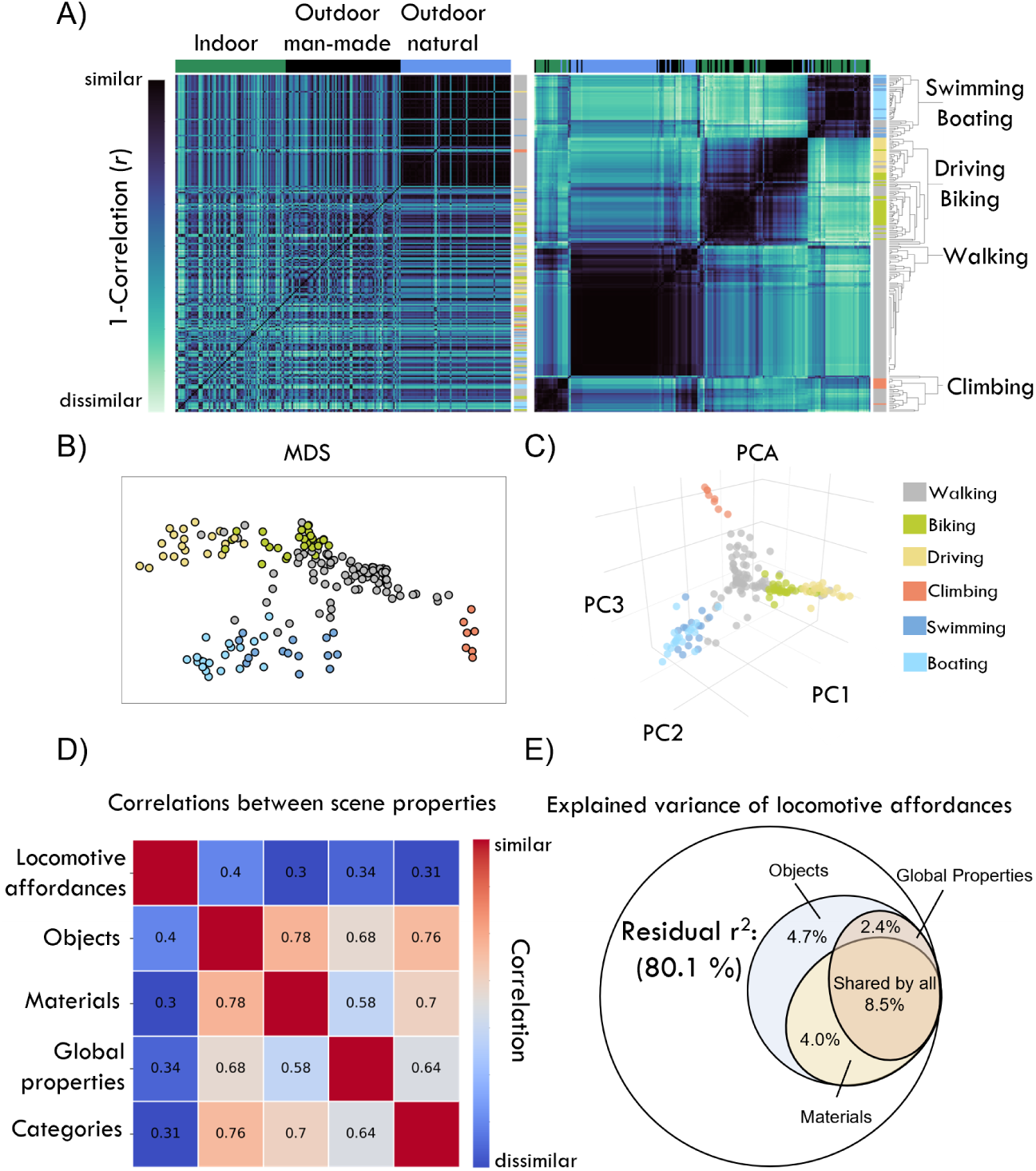
Locomotive action affordances form a distinct representational space. **(A)** RDM derived from locomotive action affordance annotations, sorted on environment type (left) or after hierarchical clustering (right). Vertical legend colors (green/blue/black) indicate different environment types. Horizontal legend colors (yellow/red/blue) indicate different actions, determined based on the predominant affordance label (see Materials and Methods). **(B)** Multi-dimensional scaling (MDS) of the action affordance RDM shows that clusters extend in three directions. A high-resolution version with image thumbnails is provided in **Supplementary Figure 1**. **(C)** Principal component analysis (PCA) of the locomotive affordance annotations reveals a three-dimensional structure with images separating along three principal components (PCs) that represent a swimming/boating dimension, a biking/driving dimension, and a climbing dimension. **(D)** Comparisons between the locomotive action affordances RDM and RDMs derived from other scene properties, indicating high correlations between objects, materials, global properties, and scene categories, and comparatively lower correlations with affordances. See **Supplementary Figure 2** for alternative RDM distance metrics. RDMs for each behavioral annotation task, as well as a comparison with automated labeling methods, are in **Supplementary Figure 3**. **(E)** Variance partitioning of locomotive action affordance annotations by the three scene properties yielding the highest combined correlation. The Euler diagram shows the unique and shared variance in locomotive affordances explained by objects, materials, and global properties. In total, 80.1% of the variance in the affordances representational space remains unexplained (residual circle; not drawn to scale).

To better understand the clustering of locomotive affordances, we visualized the dissimilarity in affordance annotations for all scenes in a 2D space by applying multidimensional scaling (MDS) to the affordance RDM (**Fig. 2B**). This shows that walking, as the most common category, forms a central hub surrounded by distinct clusters for the other actions: one extending towards biking and driving, another towards swimming and boating, and a third towards climbing. This three-dimensional structure is also evident when applying principal component analysis, a dimensionality reduction technique that computes directions in representational space that best describes the variance in affordance annotations. Visualizing the scene images along the first three principal components (PCs) shows clear separation of the locomotive affordances along these dimensions (**Fig. 2C**). These results suggest that the locomotive action affordance labels provided by our human participants form a structured representational space spanning at least three distinct dimensions.

We quantified the relation between locomotive action affordances and other scene properties by correlating the affordance RDM with RDMs derived from annotations of objects, materials, scene categories and global properties, obtained in the same behavioral experiment (**Fig. 1B**, right). Affordances show lower correlations with other scene properties (average *ρ* = 0.34, SD = 0.04) compared to how correlated these are with one another (average *ρ* = 0.69, SD = 0.07) (**Fig. 2D**). This pattern was robust across multiple distance metrics for quantifying representational dissimilarity (**Supplementary Figure 2**), suggesting affordances show minimal overlap with any of the other scene properties when considered in isolation. To test whether a combination of these properties can predict affordance perception, we conducted a variance partitioning analysis, which computes a series of linear regression models to estimate the degree of unique and shared variance between the affordance RDM and combinations of the three best correlating RDMs of other scene properties (see Materials and Methods). This analysis shows that even when combined, other scene properties only account for a small amount (19.9%) of the total variance in locomotive action affordances annotations, with the only unique contribution coming from objects (unique r^2 =^ 4.7%). Materials and objects share 4.0% of the variance in affordance annotations, global properties and objects share 2.4%, and most of the explained variance (8.5%) is shared by all three scene properties (**Fig. 2E**). This high proportion of shared variance in objects, materials and global properties highlights a well-known property of natural images, namely that they exhibit inherent covariance of visual scene properties (Malcolm et al., 2016). Therefore, our observation that locomotive action affordance annotations are not trivially predicted by (a linear combination) of other scene properties suggests they form a distinct representational space.

The limited correspondence between locomotive action affordance representations and other scene properties is surprising given our *a priori* intuition that we use (a combination of) readily available scene features to compute affordances, as well as the clear separation of action affordance annotations into distinct clusters separating, for example, water-based actions (swimming, boating) from other locomotive actions (**Fig. 2B,C**). One possibility is that our annotations of other scene properties, consisting of a handful of labels in each task, did not exhaustively capture the representation of locomotive action affordances. To test this, we also computed RDMs based on outputs of machine learning classifiers trained on the SUN Attribute Database (Patterson et al., 2014), ADE20K (Zhou et al., 2017) and Places365 (Zhou et al., 2018), which have larger sets of labels of materials, global properties, categories, and objects (e.g., 150 objects in ADE20k; 37 materials in SUN Attribute Database; 365 scene categories in Places365). Despite constituting a more exhaustive sampling, these automated labels yielded similarly low predictions of our locomotive action affordance annotations (average *ρ* = 0.21, SD = 0.09; **Supplementary Figure 3**). Moreover, classifier outputs of function labels from the SUN Attribute Database, which include several locomotive actions such as cycling, and driving, also correlated comparatively low with the same classifier’s outputs of other scene properties (average *ρ* = 0.28, SD = 0.14). These results suggest that the low predictability of human locomotive action affordance perception by other scene properties is not driven by our precise choice of annotation labels, but rather reflects a persistent difference in representational spaces.

Together, these behavioral results suggest that our selection of six locomotive action affordances of visual scenes form a three-dimensional representational space that is readily and robustly accessed by human participants, yet weakly correlates with environment types and other scene properties.

### Scene-selective PPA and OPA uniquely represents locomotive action affordances

Above, we used behavioral annotations to establish the distinct representational space of affordances for six ways people can move in the world. Next, we tested our hypothesis that these locomotive action affordances are represented in the human visual system. Brain activity was measured using a 3T MRI scanner while participants (*n* = 20) viewed a strategically sampled subset of 90 images that maintained the three-dimensional structure of the behavioral affordance representational space (see Materials and Methods and **Supplementary Figure 4**). To examine a potential influence of task instructions on affordance representation in the brain, all participants performed three distinct tasks on each of the 90 images whilst in the scanner: affordance annotation, object annotation, and an orthogonal task at fixation (see Material and Methods).

We first established that our behavioral tasks transferred well to the scanner environment by analyzing the behavior of the fMRI participants on the affordance and object annotation tasks. Average RDMs derived from in-scanner task responses (**Fig. 3A**) correlated strongly with those derived from annotations obtained in the behavioral experiment (affordances: *ρ* = 0.85; objects: *ρ* = 0.82), with a similar modest correlation between affordances and objects as in the online experiment (*ρ* = 0.44; **Supplementary Figure 5A**). In-scanner affordance annotations also exhibited reasonable agreement among subjects (mean *ρ* = 0.52; **Supplementary Figure 5B**), showing that action affordance representations are stable across both testing environments and participants, even more so than object annotations (mean *ρ* = 0.33). Overall, the in-scanner behavior replicates the original behavioral task for a smaller set of images and response options, ensuring that participants performed the same behavior while we measured activity in their visual cortex.

**Figure 3:**
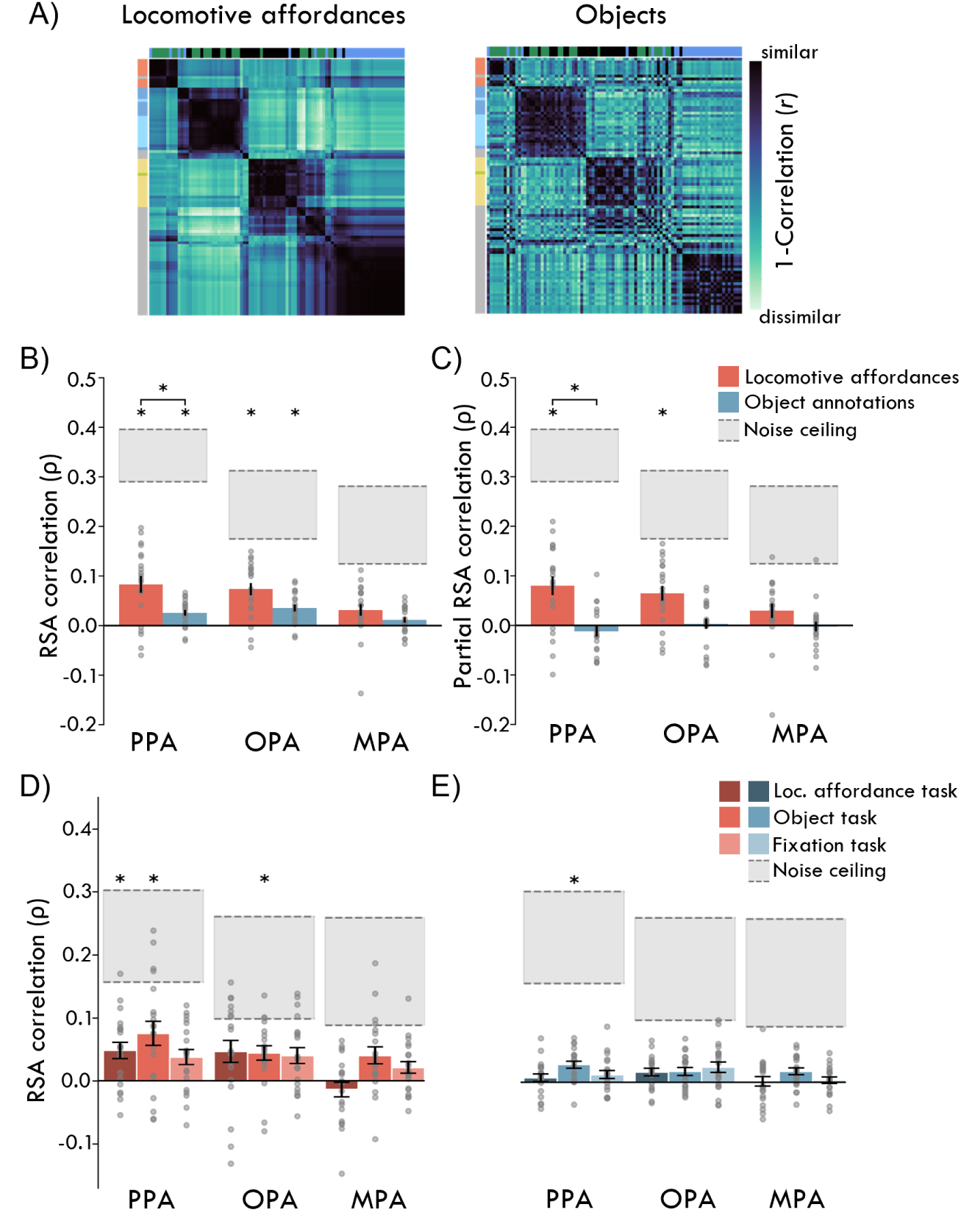
Scene-selective regions represent locomotive action affordances. **(A)** RDMs derived from in-scanner locomotive action affordance (left) and object (right) annotations, sorted by hierarchical clustering of the action affordance RDM. **(B)** Correlation of response patterns in scene-selective brain regions with locomotive action affordance (red) and object (blue) RDMs. Bars represent averages for individual participant’s (gray dots). Error bars indicate standard error of the mean (SEM) across participants. Shaded areas delineate upper and lower noise ceilings, reflecting similarity between individual participants RDMs and the group mean RDM. Asterisks indicate one-sample t-tests against zero (p*<*0.05); horizontal brackets indicate significant differences between affordance and object spaces as revealed by *P<*0.05 in paired-sample t-tests. Significance was corrected for multiple comparisons by applying Bonferroni correction across the two behavioral spaces, the three scene-selective ROIs, and both one-sample and pairwise tests. **(C)** Correlation between action affordance and fMRI RDMs, when partialling out the object space, and vice versa. Plot elements as in (B). **(D)** Task-specific correlations with the locomotive action affordances RDM (dark shades: locomotive action affordance task; medium shades: object task; light shades: fixation task). **(E)** Task-specific correlations with object annotations RDM. Shades as in (D).

To investigate what scene properties are represented in scene-selective brain areas, we first constructed RDMs based on pairwise correlation distances of multi-voxel activity patterns in each individual subject’s PPA, OPA and MPA (see **Supplementary Figure 6**), averaged across all tasks, and then correlated these RDMs with the average RDMs derived from the in-scanner affordance and object behavioral annotations, respectively (**Fig. 3A**). We found significant evidence for locomotive affordance representations in PPA and OPA (both *ρ >*0.07, *t* (19) *<*4.81, *p<*0.001) but not MPA (*ρ* = 0.03, *t* (19) = 2.59, *p* = 0.018; with all tests Bonferroni-corrected for multiple comparisons across behavioral RDMs and fMRI ROIs (**Fig. 3B**). Both PPA and OPA, but not MPA also represented object information; furthermore, through the use of paired samples t-tests, we established that responses in PPA correlated significantly higher with locomotive affordances than with objects (see **Supplementary Table 7** for full statistical summary). In contrast, OPA and MPA did not show significant differences between the two behavioral spaces. Similar results were obtained when comparing fMRI responses to individual participant behavior (**Supplementary Figure 7A**).

The significant and stronger correlation with action affordance representations compared to objects suggests that locomotive action affordances are indeed represented in scene-selective cortex. However, given the non-zero correlation between the affordance and object annotation-derived RDMs, some of the variance in fMRI response patterns may be equally well described by both spaces. To determine the degree of unique representation of affordances compared to objects, we computed partial correlations (**Fig. 3C**; see **Supplementary Table 8** for statistical summary) and found significant unique correlations with the affordance space in PPA and OPA, but not MPA, and no significant partial correlation with the object space in any ROI. Paired t-tests furthermore showed that the partial correlation with the affordance space is significantly higher than the object space for PPA, but not for OPA and MPA. These results suggest that PPA and OPA, but not MPA, represent locomotive action affordances of real-world scenes independent of their contained objects.

Comparisons of the fMRI response patterns with all five representational spaces derived from the original behavioral experiment (**Fig. 1B**) furthermore show that affordance annotations exhibit the highest correlation in all three ROIs, followed by global properties (**Supplementary Figure 7B**). Unlike objects, affordances and global properties *both* exhibit a unique correlation with PPA and OPA (**Supplementary Figure 7C**). We furthermore find that the locomotive action affordance representation in PPA and OPA is independent from broad superordinate environment type distinctions (indoor/outdoor, man-made/natural) as well as low-level visual representations captured by the GIST model, a feature descriptor used in computer vision to represent the global structural characteristics of an image, by summarizing the spectral energy across different scales, orientations and image locations (Oliva and Torralba, 2001) (**Supplementary Figure 8**). Overall, these results show that scene-selective PPA and OPA represent different locomotive action affordances in scenes, and that these representations are at least partly independent from other scene properties.

### Task-independent representation of locomotive affordances in PPA and OPA

Having established unique locomotive action affordance representations in PPA and OPA, we next asked how task instructions affect the strength of these representations in the brain. We generated RDMs based on fMRI responses separately for the three tasks performed in the scanner (affordance annotation, object annotation, fixation task) and again correlated each of these with the RDMs derived from in-scanner behavior (**Fig. 3A**). If locomotive affordances are only represented in the brain when task-relevant, we expect to find significant correlations with the affordance RDM only during the affordance task, but not the object task or the fixation task. Alternatively, if locomotive affordances are computed automatically, significant correlations with the affordance RDM are expected in all three tasks. A third possibility is that locomotive action affordances are represented when participants actively respond to the scene images (i.e., during either the affordance or object task), but not when they respond to the fixation cross (fixation task).

The results, shown in **Fig. 3D** (see **Supplementary Table 9** for statistical summary), are most consistent with task-independent computation of locomotive action affordances in PPA and OPA. PPA exhibited significant correlations with the affordance space across both the affordance and the object annotation tasks, while OPA only shows a significant correlation in the object task. However, correlations are overall of similar magnitude across tasks, and pairwise comparisons between tasks were non-significant, indicating a lack of a clear, robust task effect. Consistent with the initial task-averaged analysis, we find no significant correlation with the affordance space for any tasks in MPA. These results suggests that the representations of locomotive action affordances within PPA and OPA, but not MPA, remain largely consistent regardless of the task at hand. Representations of objects (**Fig. 3E**; see **Supplementary Table 10** for statistical summary) also do not show task-dependent correlations, with the only significant correlation found in PPA during the object task. Similar results were obtained when comparing partial correlations across tasks (**Supplementary Figure 9**). Together, these results suggest that representations of locomotive action affordances are automatically extracted by scene-selective regions OPA and PPA.

### Representation of locomotive affordances extends into mid-level visual regions

Thus far, our findings demonstrate that locomotive action affordances are represented within scene-selective ROIs. To explore whether affordance representations extend beyond these predefined regions, we performed a series of whole-brain searchlight analyses on the fMRI data. Using spherical ROIs with a 5-mm radius centered around each voxel, we extracted multi-voxel patterns spanning the entire brain volume of each participant (see Materials and Methods). Analogous to our ROI analysis, we correlated those multi-voxel patterns to our behavioral spaces derived from in-scanner affordance or object annotations (**Fig. 3A**).

The searchlight results were consistent with the ROI analysis. Significant correlations with locomotive affordances were found in OPA and PPA (**Fig. 4A**), but not in MPA. These correlations also extended beyond the scene-selective regions into other parts of visual cortex. For visualization, we overlaid the ROIs from the Julian atlas (Julian et al., 2012) that were used in our main analyses, as well as an additional ROI of early visual area V1 from the Juelich Brain atlas (Amunts et al., 2020). To account for small scene-selective regions in the Julian atlas (which includes only voxels that systematically overlap many participants) we also overlaid averaged scene-selective ROIs from a publicly available, high-powered fMRI study (the Natural Scene Dataset (Allen et al., 2022)). Comparison of these ROIs with the searchlight correlations shows that the representation of locomotive affordances extends mainly into mid-level and early visual cortex, including V1, whilst not showing any extensions into more anterior regions (e.g. parietal or premotor areas).

**Figure 4:**
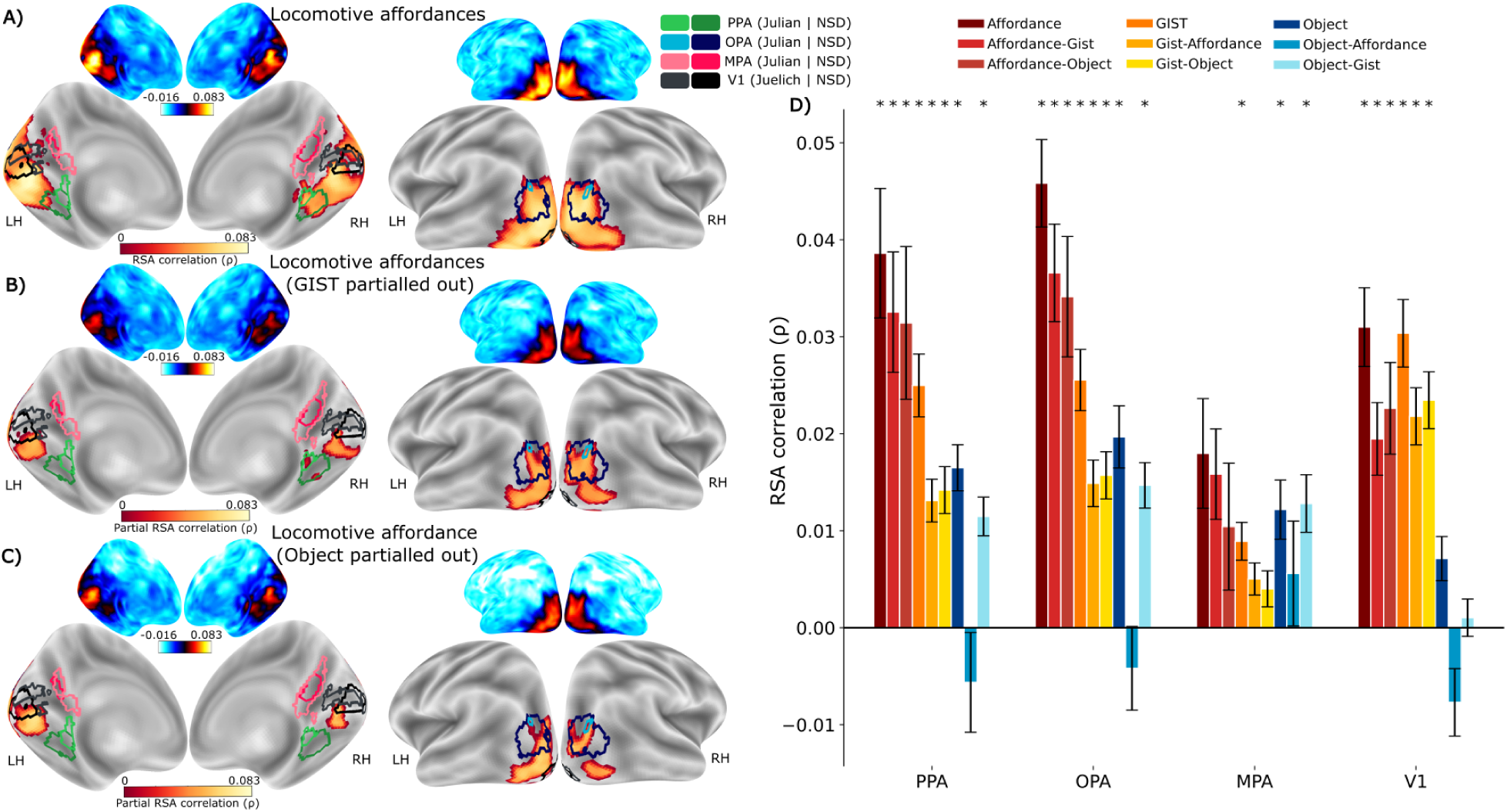
Affordance representations extend into mid-level visual regions. **(A)** Medial (left) and lateral (right) views of the average searchlight RSA correlation (Spearman’s *ρ*) with locomotive action affordances (one-sample test against zero, thresholded using a whole-brain permutation-based multiple comparison correction at p*<*0.05). Unthresholded correlation maps are shown as smaller insets. Group-level ROIs—PPA, OPA, and MPA—derived from the Julian atlas are highlighted in lighter colors and extended with the V1 ROI from the Juelich atlas. Corresponding group-level ROIs from NSD are overlaid in darker shades of the same colors. **(B)** Average partial searchlight RSA correlation with locomotive action affordances, controlling for GIST representations. **(C)** Average partial searchlight RSA correlation with locomotive action affordances, controlling for object representations. **(D)** Average searchlight correlations in predefined ROIs. Error bars indicate SEM across participants. Asterisks indicate one-sample t-tests against zero (p*<*0.05) corrected for multiple comparisons using Bonferroni correction.

Analogous to the ROI analysis, we conducted partial correlation analyses to investigate to what extent these findings can be explained by overlap with other scene properties, specifically object and GIST features. **Fig. 4B, C** highlight that accounting for this overlap reduces the extent of significant searchlight correlations, especially in early visual cortex, but does not completely eradicate correlations in mid-level visual regions. Applying the Julian atlas ROIs to the searchlight correlation maps (**Fig. 4D**) shows that the searchlight analysis replicates our ROI findings, demonstrating significant locomotive action affordance representation in the OPA and PPA, but not MPA. Notably, while significant correlations are also found in V1, this regions shows overall lower correlations compared to PPA and OPA, with a comparable correlation for affordances as for GIST. Furthermore, controlling for overlapping features by partialling out the GIST or object features substantially reduces the correlations with locomotive affordances in this area.

In sum, searchlight analyses reproduce our hypothesis-driven ROI analysis, providing robust evidence for locomotive affordance representations in scene-selective regions, while also demonstrating extensions into lower and mid-level visual regions.

### Pre-trained DNNs show weak alignment with locomotive action affordances

To further investigate the representations underlying locomotive action affordance perception, we again used RSA to examine to what extent deep neural networks (DNNs) can capture human behavioral and neural responses to our image set. We sample models with different architectures (both CNNs and Transformers), task objectives (object/scene classification, scene segmentation, video classification) and training procedures (supervised learning using labels, contrastive learning with image-text pairs, self-supervised learning) and focus on the comparative ability of these models to capture the similarity between human annotations of perceived locomotive affordances versus objects. We computed RDMs from the DNN activations to the scene stimuli in a subset of layers in each model (see Materials and Methods), and report the layers that correlate the highest with human behavior or brain responses (correlations for all individual layers of a few example models are provided in **Supplementary Figure 12**).

All DNNs demonstrate significant correlations with both locomotive action affordance and object representations (all *t >*17.87, all *p <*0.001; **Fig. 5A-B**). However, across all models, correlations were substantially lower with locomotive action affordances (average *ρ* = 0.19, SD = 0.06) than with objects (average *ρ* = 0.31, SD = 0.07;pairwise *t* (12) = 10.23, *p <*0.001). Classic CNNs optimized for object classification on ImageNet correlated the least with human behavioral action affordances, but correlations increased for CNNs trained on scene datasets, with the highest correlation for a CNN trained on semantic segmentation (ResNet50 trained on ADE20k: *ρ* = 0.21). Video-trained CNNs trained on action recognition (X3D and SlowFast) and CNNs trained with alternative training objectives (DINO or CLIP) show comparable correlations as these scene-dataset trained models, while Vision Transformers (ViTs) exhibit the highest correlations with locomotive affordances (maximum for ViT Base Patch 16 with *ρ* = 0.29). Human object annotations (**Fig. 5B**) also showed the highest correlation with the ViT Base Patch 16 (*ρ* = 0.38).

**Figure 5:**
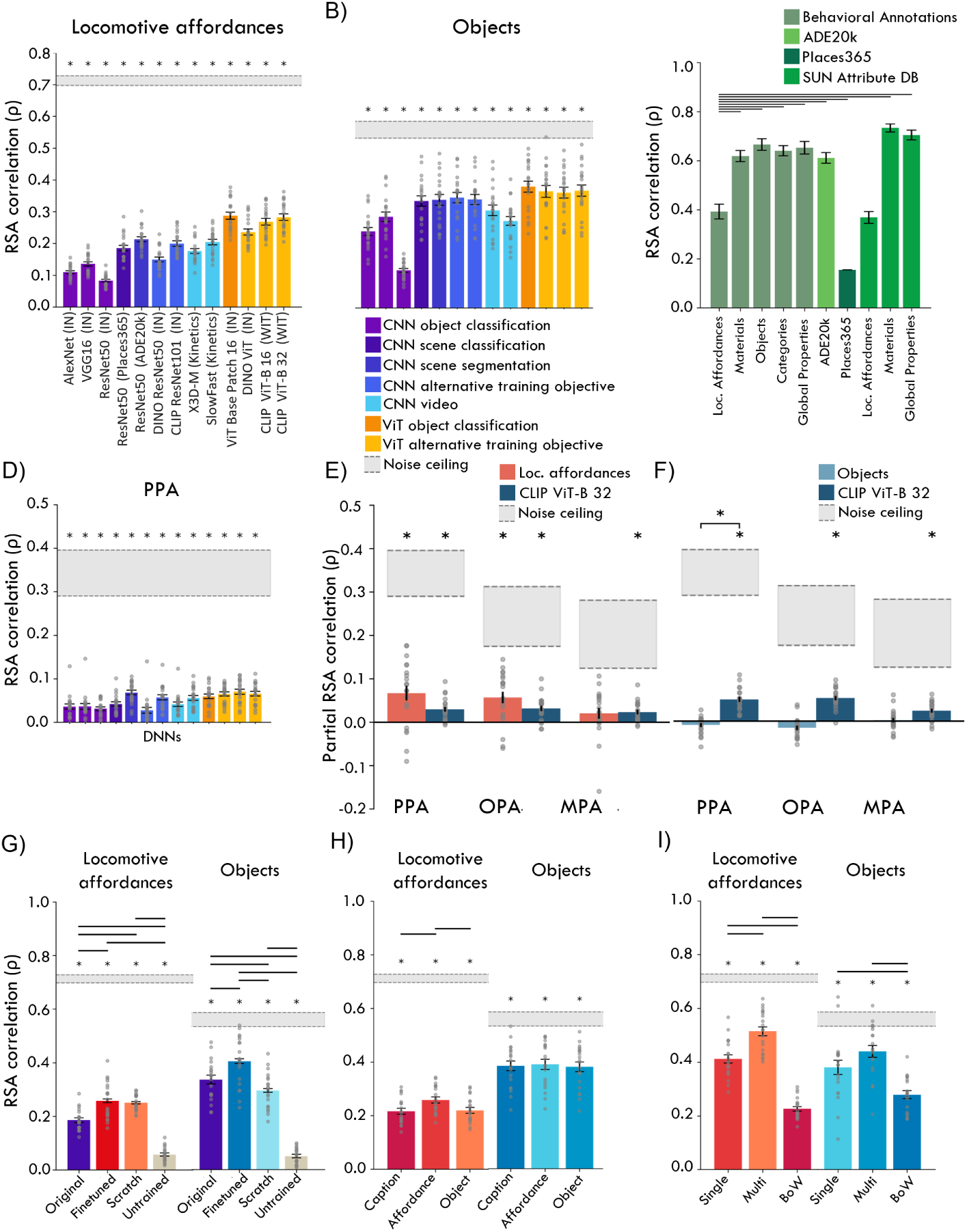
Comparison of DNNs with human behavior and fMRI responses in scene-selective cortex. **(A)** Correlation between in-scanner locomotive action affordance annotation RDMs and RDMs derived from feature activations from DNNs with different architectures and training objectives. Bars represent average across participants for the best correlating layer (see **Supplementary Table 1**); gray dots indicate individual participants. Significant t-tests correlations are marked by asterisks indicating p*<*0.05 corrected for multiple comparisons using Bonferroni correction. The shaded area delineates upper and lower noise ceiling; error bars indicate SEM across participants. **(B)** Correlation of DNN RDMs with in-scanner object annotation RDMs. Plot elements as in (A). **(C)** Correlation of ViT Base Patch 16 with annotations for the full set of images from the online behavioral experiment and automatic labeling outputs. Black horizontal lines indicate significant (p *<*0.05) pairwise comparisons with a two-tailed sign test (FDR-corrected). Error bars indicate 95% confidence intervals derived from a bootstrap distribution (n = 1000). **(D)** Correlation of each DNN’s best correlating layer with PPA (averaged across tasks); OPA and MPA are reported in **Supplementary Figure 10**. Plot elements as in (A). **(E)** Correlations between scene-selective cortex and locomotive action affordances, when partialling out CLIP ViT-32. **(F)** Same as E but for objects. **(G)** Correlation between in-scanner human behavior and four different ResNet50 models: original (pretrained on Places365, same as in A-B), finetuned (last fully connected layer retrained on function labels of Scene Attribute Database), scratch (trained end-to-end using function labels), untrained (randomly initialized). **(H)** Correlations between in-scanner human behavior and RDMs derived from LLM embeddings of three types of image captions (general caption; action affordance-centered caption, object-centered caption; see **Supplementary Table 2**. **(I)** Correlation between in-scanner human behavior and RDMs derived from GPT-4 outputs, using different annotation prompts: choose a single (Single) or multiple (Multi) affordances/objects from the same options human participants had; or list a maximum of 50 possible actions/objects in the scene (BoW; see **Supplementary Table 3**). (G-I) plot elements as in (A).

These results suggest that, unlike human PPA and OPA, the DNNs we tested generally exhibit higher correspondence with object representations than with locomotive action affordances. Indeed, even the best correlating DNN with in-scanner affordance annotations (ViT Base Patch 16) still correlated significantly higher with objects [pairwise *t* (19) = 3.66, *p* = 0.0017]. Moreover, the relatively large gap between the best-correlating model and the noise ceiling for locomotive affordance behavior shows that a substantial amount of variance in the stable affordance representational space in humans is unexplained by DNNs. We find the same pattern when comparing this ViT with the behavioral annotations obtained in the original behavioral experiment for the full set of images, as well as automated labels obtained with the scene segmentation model trained on ADE20k, scene categories from Places365 and attribute classifiers (**Fig. 5C**): locomotive affordance annotations, whether obtained from human behavioral annotations or automatic labeling, show significantly lower correlations with the ViT activations than with all other representational spaces (with the exception of the sparse scene categories from Places365), again demonstrating their distinct representation.

Together, these results demonstrate significantly lower alignment of DNNs with locomotive action affordances compared to objects. While this is unsurprising for DNNs explicitly trained to classify objects, we note that this result also holds for models trained on other tasks such as scene classification and scene segmentation, and models trained with self-supervised (DINO) and contrastive learning objectives (CLIP).

### Affordance representation in PPA and OPA is independent of DNN features

To determine how well the different types of DNNs align with human brain responses to our novel stimulus set, we also compared their feature activations with fMRI response patterns in PPA (**Fig. 5D**), and OPA and MPA (**Supplementary Figure 10**). All models show significant correlations with the scene ROIs (PPA: all *t* (19)*>*4.08, *p<*0.001, OPA: all *t* (19)*>*4.51, *p<*0.001 and MPA: all *t* (19)*>*4.71, *p<*0.001). Mirroring behavior, ViTs showing numerically highest correlations in all three ROIs (PPA v. CLIP ViT-B 32, *ρ* = 0.070; OPA v. CLIP ViT-B 32, *ρ* = 0.061; MPA v. CLIP ViT-B 16, *ρ* = 0.043) although the ResNet50 trained on scene segmentation performs on par (PPA, *ρ* = 0.068). However, model correlations are substantially below noise ceiling, and models show less differentiation here compared to behavioral measurements (**Fig. 5A-B**).

Partial correlation analyses furthermore show that the correlations of DNN features with fMRI responses in scene-selective cortex primarily reflects object-related information. Both PPA and OPA exhibit significant unique correlations with locomotive affordances when partialling out the best correlating DNN (CLIP ViT-B 32), while the DNN itself also explains unique variance in all three ROIs, **Fig. 5E**; see **Supplementary Table 11** for statistical summary). In contrast, neither PPA, OPA or MPA show significant positive correlations with object annotations when partialling out DNN features (**Fig. 5F**), whilst showing significant unique correlations with the DNN features. These results suggest that DNN features do not fully account for locomotive action affordance representation in PPA and OPA, instead capturing object representations in these regions.

### Enhancing alignment with human affordance perception using supervision and LLMs

Our results point to a consistent gap in DNN’s abilities to explain locomotive affordance compared to object representations reflected in human behavioral and brain measurements. In this final section, we explore three ways in which DNN alignment with human affordance perception could potentially be improved.

First, while we tested a variety of DNNs, none of them was *explicitly* trained to perform locomotive affordance perception. One approach towards closing the gap between DNN object vs. affordance perception could be to simply train the model on affordance labels, instead of object labels. We therefore assessed if DNN alignment with human behavior increases with explicit supervision with affordance labels, using the SUN Attribute Database subset of function labels (see Materials & Methods and **Supplementary Table 4**). We find that both end-to-end training and fine-tuning on these labels improves correlations with human-perceived affordances, relative to training on scene classification only (**Fig. 5G**) (fine-tuned vs. original Δ*ρ* = 0.07, pair-wise t-test *t* (19) = 6.89, *p<*0.001; end-to-end training vs. original Δ*ρ* = 0.06, *t* (19) = 7.5, *p<*0.001). This shows that with direct mapping to affordance labels, visual features extracted from (pre-trained) DNNs can better approximate human-perceived affordances. However, even with such direct supervision, the gap in alignment relative to objects remains (average difference in *ρ* = 0.15), and the obtained increase due to training on affordance labels is relatively modest, remaining lower than the best performing ViT model trained on object recognition (**Fig. 5A**).

Second, prior work has proposed that human vision is not only shaped by visual features but also by language (e.g. Simanova et al., 2016). For example, similarity judgments of visual stimuli are not only captured by visual features, but also by linguistic description such as captions (Marjieh et al., 2022). Moreover, recent fMRI studies suggest that DNNs trained with richer linguistic descriptions yield better predictions of high-level visual cortex responses to natural scenes (Wang et al., 2023; Doerig et al., 2022). Consistently, our results indicate that visual features extracted from CLIP models - trained to pair images with captions - are among the best ranking models in terms of alignment with human behavior and fMRI responses (**Fig. 5AB,D**), although notably they do not show substantially higher correlations than the ViT Base Patch 16 or the scene segmentation model, which were not trained with rich linguistic descriptors. To examine to what extent linguistic representations capture locomotive action affordance perception, we used a multi-modal large language model (GPT-4, OpenAI et al. (2023)) to generate an affordance-centered, an object-centered caption, and a general caption for each image in our stimulus set (see **Supplementary Table 2** for examples). These captions were then fed to an LLM (see Materials and Methods) to extract language embeddings, from which we computed RDMs to compare to our human behavioral annotation RDMs.

As shown in **Fig. 5H**, LLM embeddings of all three types of captions correlate significantly with the human perceived locomotive affordances (general caption *ρ* = 0.22; affordance caption *ρ* = 0.26; object caption *ρ* = 0.22) all *t* (19)*>*19.65, *p<*0.001). LLM embeddings of affordance-centered captions correlate significantly higher with human locomotive action affordance behavior than the other two types of captions (both pairwise *t* (19)*>*8.02, *p <*0.001). However, all three LLM embeddings still show higher correlations with object annotations (general caption *ρ* = 0.38, affordance caption *ρ* = 0.39, object caption *ρ* = 0.38; all *t* (19)*>*20.54, *p<*0.001), with no significant pairwise differences. This indicates that while LLMs capture scene semantics to some extent, they still exhibit an alignment bias towards visual object representations versus locomotive action affordances, similar to DNNs. Moreover, we find that the LLM-based captions do not capture fMRI responses in PPA and OPA well, showing substantially lower correlations compared to the affordance space, and no significant partial correlations (see **Supplementary Figure 13**).

Finally, we explored if it is possible to use the multi-modal LLM to predict locomotive affordances directly, by conducting an analogous behavioral experiment with GPT-4 as with our human participants. We presented our images to GPT-4 with a few different prompts, asking it to select one (Single) or multiple (Multi) actions/objects from the same list of response options as given to our human participants, or to name all possible actions and objects in the image (BoW; see **Supplementary Table 3**). We then created RDMs by converting the model outputs to one-hot vectors (see Materials and Methods) and correlated them to the RDMs derived from human behavior. As shown in **Fig. 5I**, GPT-4 generates behavioral output that now aligns to a similar extent with locomotive affordances as with object annotations, with overall highest correlations when given the exact same task as the participants (affordances: *ρ* = 0.515; objects:*ρ* = 0.440). Notably, however, even with the identical annotation task, a gap relative to the noise ceiling still remains, suggesting GPT-4 does not fully emulate human behavioral annotations.

These findings suggest that DNNs trained on object or scene classification, as well as several other commonly-used visual understanding tasks, capture only some aspects of human scene representations. Specifically, we show that good alignment of visual DNN features with human object representations does not automatically transfer into good alignment with affordance representations. Training DNNs directly on affordance labels increases alignment but does not fully close the gap with object recognition; while a multi-modal LLM can partially recapitulate human affordance annotations, LLM-generated linguistic descriptions alone show comparatively poor alignment with human affordance representations. Collectively, our comparisons thus suggest that the networks we tested do not fully capture locomotive action affordance perception in humans.

## Discussion

The aim of our study was to assess whether the human visual system represents different locomotive actions of complex natural scenes, and if these representations are distinct from other scene properties. The notion of action affordances shaping visual perception is long-standing in psychology (Gibson, 1977), but the presence of distinct action affordance representations for scene environments in the human brain had so far not been demonstrated. Utilizing behavioral and fMRI measurements alongside computational deep neural network models, we discovered that human perceived locomotive affordances - reflecting different ways we can move through our local environment - form a unique representational space that is not trivially predicted from other perceived properties such as objects or global layout. Moreover, scenes associated with specific locomotive actions, such as driving or swimming, evoked similar multi-voxel activation patterns in PPA and OPA (but not MPA), with representations extending into early- and mid-level visual regions. By accounting for variation in other visual scene properties, we find that these locomotive action affordances are indeed represented in the brain independent from objects and other (low-level) scene properties. Importantly, these representations proved to be task-independent, suggesting that OPA and PPA encode distinct locomotive action affordances automatically even when participants passively view the images or perform a non-navigational task. While pre-trained DNNs partially explained activity in scene-selective regions, their alignment with locomotive affordances was lower than with object representations, suggesting that such features do not fully account for human affordance perception.

Prior fMRI research on affordance perception has primarily focused on the representation of navigational affordances, with evidence accumulating that OPA in particular encodes egocentric information relevant to navigate the immediate environment (Persichetti and Dilks, 2016; Dilks et al., 2011; Julian et al., 2016; Kamps et al., 2016b,a). In these studies, affordances are typically operationalized as the presence or absence of ’navigable space’ in a scene, for example in the form of walkable pathways within images of indoor rooms (Bonner and Epstein, 2017), or distance to navigational boundaries in hallways (Park and Park, 2020). In contrast, our study operationalizes affordances as multiple possible locomotive action labels in both indoor and outdoor scenes. By considering multiple types of environments and navigational actions other than walking, our study therefore extends these prior findings by suggesting that OPA may additionally represent *different types* of locomotive actions that one could perform in navigable space. A recent study by Jones et al. (2023) also investigated if OPA differentiated other locomotive actions than walking, specifically crawling and flying - neither of which were included in our behavioral annotations - and found selective responses to videos taken only during walking. This could mean that neural representation of distinct action affordances may be restricted to a relatively narrow range of locomotive actions, although methodological differences (the use of videos and analysis of univariate rather than multivariate responses) could also explain the discrepancy with our study.

In addition to OPA, we here also find representation of locomotive action affordances in PPA, which has been previously shown to represent a wide range of scene properties, including scene category (Walther et al., 2009), and spatial layout (Epstein and Kanwisher, 1998; Kravitz et al., 2011; Park et al., 2011), but also textures (Cant and Xu, 2012; Henriksson et al., 2019) and objects (Janzen and van Turennout, 2004; Harel et al., 2013; Marchette et al., 2015). In fact, the wide range of proposed roles of PPA in visual scene representation (see Epstein and Baker, 2019; Bartnik and Groen, 2023, for recent reviews) and the inherent correlations between different scene properties (Malcolm et al., 2016; Greene and Hansen, 2020; Lescroart et al., 2015) motivated us to collect multiple types of annotations for our scenes, in order to directly compare these against our new operationalization of action affordance representation in scene-selective cortex. As shown in **Supplementary Figure 7B**, we find that both locomotive action affordances and global properties are consistently represented in PPA, while scene categories and objects are less consistently represented. Importantly, representations of global properties do not ’explain away’ affordances in PPA (or OPA) (**Supplementary Figure 7B**), and results were robust across multiple ways of assessing affordance representation (in-scanner behavior, separate online experiment, and automatic labeling using classifiers). In contrast, the lack of representation of locomotive action affordances in MPA is consistent with a substantial body of work showing that MPA is involved in navigating the broader, rather than the immediate environment (Epstein and Vass, 2014; Park and Chun, 2009), linking visual inputs to cognitive maps in hippocampal systems (Vann et al., 2009; Dilks et al., 2022).

In addition to expanding the operationalization of affordances to different types of locomotive actions, we also assessed the task-dependency of these representations by measuring brain responses to our scenes under three different task instructions. Prior evidence on task effects on occipitotemporal cortex representations is mixed, with some studies finding modulations of representational patterns depending on in-scanner tasks in object-selective (Harel et al., 2012; Hebart et al., 2018) and scene-selective cortex (Lowe et al., 2016; Groen et al., 2018; Bracci et al., 2017), while others observed representations of spatial layout in PPA (Kravitz et al., 2011) or pathways in OPA (Bonner and Epstein, 2017) even when not task-relevant. A direct comparison between a categorization and navigation task on neural representations of indoor walkways found task-specific effects in PPA, but not OPA (Persichetti and Dilks, 2018). Here, we find that both OPA and PPA show task-independent representations of action affordances, while object representations in these regions were more task-dependent. Task-independent representations of locomotive affordances in scene-selective cortex are consistent with classic psychophysical findings by Greene and Oliva (2009) highlighting visual properties relevant to navigation are processed within very brief glimpses, as well as M/EEG studies showing early modulation of neural responses by scene properties (Groen et al., 2013; Harel et al., 2016), including affordance-related information (Harel et al., 2022; Djebbara et al., 2021). The results of our searchlight analysis also highlight that the representation of locomotive affordances may involve relatively early visual responses. It is possible that more task-dependent representations are present in other brain regions, including dorsal stream parietal cortex or frontal cortex (Bracci et al., 2017; McKee et al., 2014).

DNNs have recently emerged as a popular tool for studying computations underlying human visual perception (Kietzmann et al., 2019; Storrs and Kriegeskorte, 2019), by mapping representations of DNN features to different brain regions (e.g., Khaligh-Razavi and Kriegeskorte, 2014; Dwivedi et al., 2021a). Studies focusing specifically on scene-selective regions found that they tend to correlate highest with mid-level layers in object- and scene-classification trained CNNs (Bonner and Epstein, 2018; Groen et al., 2018; King et al., 2019); here, we find the same pattern for these CNNs, but models with other architectures or training objectives show a more complex pattern (**Supplementary Figure 12**). Furthermore, Dwivedi et al. (2021b) pointed out that DNNs trained on scene segmentation, i.e. with the objective to spatially locate different elements of a scene, show a higher alignment with scene-selective regions; we observe a similar advantage in our data (**Figure 5D**). More in-depth inspection of the model features driving DNN alignment with scene-selective regions in (Bonner and Epstein, 2017) showed that it is driven by presence of extended surfaces and floor elements in the lower visual field (Bonner and Epstein, 2018; Dwivedi et al., 2021b), consistent with the general notion that these regions contain visuospatial representations (Groen et al., 2022).

While we here also find significant correlations of DNN features with scene regions, they do not correlate strongly with behavioral locomotive action affordance annotations, and do not explain ’away’ the variance in the brain response related to affordances (**Figure 5E**). Instead, we find that all DNNs are more closely aligned with object representations - this distinction was consistently observed across multiple DNN architectures trained on a variety of tasks and datasets, arguing against the idea that affordance representations emerge ’automatically’ from DNN features learned for the purpose of image classification (Bonner and Epstein, 2018). The relatively poor alignment of DNN features with our behavioral and brain responses may reflect our operationalization of affordances through action labels, which are potentially less spatial and more semantic in nature than navigable pathways. However, our findings do not necessarily imply that locomotive affordance representations are not linked to visual features in scenes at all: our DNN training and fine-tuning results (**Figure 5G**) show that it is possible to achieve enhanced prediction of behaviorally perceived affordances using direct supervision with affordance labels, suggesting that relevant image features can be learned by DNNs, but that they might be different from those useful for object recognition.

We also explored whether non-visual, linguistic features could explain the gap in DNN alignment with affordance perception. Our results show that purely linguistic descriptions of scenes, even when containing affordance-related information, only partially close this cap, still showing lower alignment with behavioral annotations of affordances than with objects (**Figure 5H**), and low correlations with scene-selective cortex responses (**Supplementary Figure 13**). In addition, visual features extracted from DNNs trained with richer linguistic supervision do not show substantially higher correlations compared to models trained with single labels (**Fig. 5AB,D**). Interestingly, we do find that the gap in alignment with affordance behavior relative to objects can be closed when using a GPT-4-vision to emulate human behavior directly (**Fig. 5I**), suggesting this particular multi-modal model has achieved a similarly efficient mapping of relevant image features for affordance labels as for object labels. This serves as a proof of principle that such a mapping can be learned, at least for the particular labels used our study. However, one notable difference between DNNs and humans that could be an important factor in explaining the misalignment is that these models lack embodiment. Indeed, the classic ecological psychology view on affordance perception states that it is fundamentally shaped by the (bodily) abilities possessed by the observer (Gibson, 1977; Rietveld and Kiverstein, 2014). Therefore, one potential promising direction towards closing the alignment gap would be to compare human perception with embodied AI systems that perform the kinds of visually guided locomotive behaviors examined here.

Several key questions remain for future work. First, we only sampled a subset of pretrained DNNs, and it is possible that a DNN we did not consider here, or a different alignment method (Sucholutsky et al., 2023) than RSA will provide better predictions of human affordance perception. Our goal here was to explore relative differences in alignment of affordances relative to other scene properties, not to provide a systematic comparison of how different DNN aspects (architectures, training datasets, task objectives) affect alignment with human perception (Conwell et al., 2022; Muttenthaler et al., 2023; Sartzetaki et al., 2024). To facilitate such a comprehensive assessment as well as further exploration of the action affordance representational space, we make our stimuli, behavioral and neural data openly available, allowing testing of more models, as well as comparison with other public fMRI and behavioral benchmarks. Second, our analyses involving fine-tuning and end-to-end training DNNs showed only modest improvements in human alignment; this gain could potentially be further improved by increasing supervision signal quality. Here, we used Sun Attribute Database labels by Patterson and Hays (2012); Patterson et al. (2014), which were not collected for the purpose of studying affordances per se, and notably do not include the label ’walking’, which as shown in **Figure 1A** strongly drives human representational similarity. A dedicated large scale-dataset of locomotive action affordance annotations is needed to precisely assess how well human affordance perception can be mimicked using DNNs trained via direct supervision. Third, our comparison of task effects may be weakened by our experimental design in which participants were only probed in a subset of trials in each fMRI run; although response probing was randomized across trials and therefore unpredictable for the participants, incentivizing them to perform the task throughout the run, stronger task effects may be observed when participants are consistently probed on each trial.

Finally, we here tested locomotive action affordance perception on static images for which participants indicated how they would move in a scene image using simple button presses. This is arguably substantially different from real-world vision, in which humans experience 1) a continuous flow of input and 2) actively navigate the world. Future work should ideally combine more continuous inputs with more immersive and realistic navigational tasks (Gregorians and Spiers, 2022; Zhang and Gallant, 2020). In terms of computational models, a promising direction to close the gap in human alignment may be to consider DNN models that are trained on visual inputs that better approximate our daily navigational experiences (Greene et al., 2024; Venkataramanan et al., 2024). Moreover, it would be very interesting to explore alignment of human affordance perception with vision-language-action models from robotics (e.g. Kim et al., 2024), as these models have been trained in what could be considered more embodied contexts. Overall, our results suggest that to understand the visuo-semantic transformations underlying human perception of scene affordances, we need to move beyond object recognition and consider a wider spectrum of ecologically relevant behaviors.

To summarize, our key finding is that locomotive action affordances are distinctly represented in behavioral and neural measurements of human visual perception. We performed extensive comparisons to demonstrate that this space can be reliably measured across different experimental paradigms and in fMRI. We also explored the relation of these representations with different types of visual information derived from human behavioral annotations, automatically derived labels using classification, feature activations in a large set of DNN models, or by training or prompting such models explicitly. Our results suggest human representations of scene affordances are not fully captured by any of these models, thus offering a new human-alignment challenge for computational models of visual processing.

## Materials and Methods

### Stimuli

We created a novel stimulus set of 231 high-resolution color photographs (1024×1024 pixels) of visual environments. To prevent overlap with DNN training sets, images were collected from Flickr (freely available under either CC0 1.0, CC BY 2.0 and CC BY-SA 4.0 licenses) rather than any open large-scale image database commonly used to train DNNs (e.g., ImageNet, Places365, MS COCO). Images portrayed typical everyday environments devoid of prominent objects, humans, or animals. Each photo was taken from a human-scale, eye-level perspective, ensuring ecological validity (refer to **Fig. 1A** for exemplars). Diverging from previous research that predominantly utilized indoor imagery (e.g., Bonner and Epstein, 2017), our stimulus set contains images from three superordinate scene classes: indoor, outdoor-natural, and outdoorman-made. This broad division is predicated on the premise that different environmental contexts afford distinct functionalities (Zhou et al., 2018). A t-distributed Stochastic Neighbor Embedding (t-SNE) visualization of our stimulus set, displayed on the attribute space of over 12,000 images from the SUN Attribute Database (Patterson and Hays, 2012) can be found in **Supplementary Fig. 11**, demonstrating that our stimulus set covers a much wider range of stimuli compared to previous fMRI studies. Images were matched in luminance to the mean across the stimulus set using the SHINE toolbox (Willenbockel et al., 2010).

### Behavioral Annotations

Humans naturally engage in a diverse range of actions within natural environments. However, the visual features within these environments often co-vary with each other, complicating the identification of the essential features for specific tasks (Malcolm et al., 2016; Greene and Hansen, 2020). To determine the relation between locomotive action affordances and various visual scene features, we ran an online study in which we collected behavioral locomotive action affordance annotations along with annotations of multiple readily identifiable scene properties: contained objects, materials, global properties, and scene category (see ’Behavioral task descriptions’ below). Within the same experimental session, participants performed two additional tasks that are not further analyzed in the current paper.

#### Subjects

The online experiment initially involved 167 participants (81 female) recruited from prolific.ac (Palan and Schitter, 2018). Of these, 15 participants did not complete the study, resulting in a final count of 152 participants. The participants had a median age of 25 years (range 18 - 74 years, SD = 14). We used selection criteria provided by Prolific to screen for reliable participants (Highest education level completed: High school diploma/A-levels; normal or corrected-to-normal vision: yes; First Language: English; Approval Rate: 90–100). On average, participants spent about 39 minutes completing the experiment (SD = 18 minutes). For their involvement in the study, each participant received a compensation of 7.5 pounds (GBP). Participants provided informed consent by selecting a checkbox on a digital consent form. The Ethical Committee of the Computer Science Department at the University of Amsterdam approved the experiment.

#### Behavioral task descriptions

The experiment was designed using the Gorilla Experiment Builder (Anwyl-Irvine et al., 2020) and began with an introductory screen that informed participants about the study, instructions on how to switch into full-screen mode, and the tasks they would undertake. The experiment featured seven distinct tasks, of which the five following tasks are included in this study. In the *Take Action* task, participants selected from six options (walking, biking, driving, swimming, boating, climbing) to indicate how they would navigate through a scene. In the *Scene objects* task participants selected from six options (Building/Wall, Tree/Plant, Road/Street, Furniture, Body of water, Rocks/Stones) the presence of certain objects in the scene. This selection of objects was inspired by the objects that can be perceived in a single fixation (Fei-Fei et al., 2007). The *Scene Materials* task involved indicating the presence of ten different materials (Water, Wood (Not tree), Vegetation, Dirt/Soil, Sand, Stone/ Concrete, Snow/Ice, Pavement, Carpet, Metal), which were inspired by the material labels used in the SUN attributes database (Patterson et al., 2014). The *Scene Categories* task asked participants to select from eight types of scene categories (Room scene, Inside city scene, Hallway scene, Forest scene, Desert scene, Open country scene, Mountain scene, Coast/River scene) which ones applied to the scene. In all of these tasks, participants were allowed to select one or more annotations without any time limit. In the task *Scene Attributes*, participants used sliders on various bipolar dimensions (Open/Closed, Man-made/Natural, Near/Far, Navigable/Non-navigable) which were inspired by prior research by Kravitz et al. (2011) and Greene and Oliva (2009). The two tasks that were not further analyzed in this paper were *Path Drawing* (involving tracing potential pathways on an image) and *Scene Description* (asking participants to encapsulate each scene in a single word using a text box).

#### Behavioral task procedure

Participants were divided into one of five groups to counterbalance the task sequence and ensure that each participant viewed each image only once in one of the tasks. In each task, members of a specific group viewed the same set of 33 images and performed the required annotations. This approach guaranteed that each image received ratings from a similar number of participants. The order of images within each task was randomized for each subject to prevent bias. After completing the seven tasks, participants filled out a feedback form with questions about their experience during the experiment. Each of the 231 real-world scene images was rated across all tasks by at least 20 participants and on average 21.7 (SD = 0.87) participants. The time taken for each annotation task was relatively consistent, with a median duration of 4.3 minutes (ranging from 3.32 to 7.95 minutes).

#### Behavioral data analysis

For the behavioral annotations, the button presses for the different tasks were recorded. Representational dissimilarity matrices (RDMs) were computed using 1-Pearson correlation distances between the proportion of participants that annotated the presence or possibility of a given label (e.g., the proportion of participants that annotated the scene as affording walking). These features were first standardized across the images by removing the mean and scaling to unit variance. Similar results also can be obtained using when not scaling or using different distance metrics (see **Supplementary Fig. 2**). This resulted in five symmetric 231×231 RDMs, one for each annotation task.

### Automatic labeling

#### Places365

To capture a richer space of possible scene categories we used a PyTorch implementation of a Wider-ResNet18 trained for scene classification on Places365 (Zhou et al., 2018) to classify each of our images to one of the 365 scene classes. This model is part of the unified code provided by CSAILVision https://github.com/CSAILVision/places365/blob/master/run_placesCNN_unified.py. The resulting one-hot vector were compared between all pairs of images to compute RDMs using Euclidean distance metric. We used Euclidean distance here since each image was assigned a single class label, rendering the computation of 1-Person correlation inapplicable.

#### SUN Attribute Database

To better understand how the choice and number of available labels affected the relation between locomotive action affordance representations and other scene properties, we utilized the Sun Attribute Database (Patterson et al., 2014). We again employed the PyTorch implementation of a Wider-ResNet18 trained for scene classification on Places365 (Zhou et al., 2018). The provided script extracts features from the model’s last convolutional layer (512 feature values) and applies a linear transformation to 102 attribute dimensions. The 102 scene attributes are grouped into 4 superordinate groups, namely functions (37 labels), materials (37 labels), spatial envelope (16 labels), and surface properties (12 labels). Notably, scene functions contain locomotive action affordance descriptions that partially overlap with the ones used in our behavioral experiment (e.g. cycling, swimming) but also more diverse activities such as socializing and gaming. Therefore, we also created an additional subspace closer to our locomotive affordance/action labels by only including the functions related to navigation (9 labels). We passed all our 231 images through the classifier and created RDMs based on a subset of the resulting attribute vector for each image standardized across the images and used 1-Pearson correlation as distance metric. This resulted in three additional RDMs representing locomotive action affordances, materials and global properties (spatial envelope).

#### ADE20K

To automatically obtain a representational space based on the co-occurrences of objects in scenes, we utilized a scene segmentation network model consisting of a ResNet50dilated encoder and a PPM-deepsup (PPM + deep supervision trick) decoder that classifies each pixel to belong to one of 150 object classes (Zhou et al., 2017). We passed the images through the network model, and extracted the unique predicted object labels for each image as one-hot vectors, e.g. if an object such as a ”wall” is present in the scene or not. Again, the feature vectors were standardized across the images and 1-Pearson correlation was used to obtain the dissimilarity in the labeled object classes resulting in one RDM representing the object/co-occurrence representational space.

### fMRI experiment

#### Stimuli

A subset of the stimuli from the online experiment were used for the fMRI study. To select a representative subset, we utilized Principal Component Analysis (PCA) on the behavioral action affordance annotations from the online experiment to extract the underlying dimensions. As walking was for most images the predominant label, we balanced the selection by labeling each image with the next most often selected locomotive action, if it was labeled by over half of the participants. Subsequently, we selected a subset of 90 images using two selection criteria. Firstly, we ensured an equal distribution of images from each of the three environmental categories – indoor, outdoor-natural, and outdoor-man-made – allocating 30 images to each type of environment. Further, we balanced the number of images for each action affordance across the environments to the maximum extent possible, selecting images with high scores on the three principal components in the locomotive action affordance PCA (**Figure 2C**). The selection of the fMRI subset relative to the full image set is depicted in **Supplementary Figure 4**.

#### Subjects

In total 20 healthy participants (13 female) with a median age of 24 years (range 18 - 33 years, SD = 3.59) completed the fMRI experiment, which consisted of four scanning sessions: an initial functional localizer session and three subsequent main experiment sessions in which participants were presented with scene stimuli under three different task instructions (see ’Experimental design’ below). Three additional participants did not complete all sessions and are therefore not included in the data analysis. All participants had normal or corrected-to-normal vision, reported no neurological disorders nor used related medication. Before participating in each session participants filled out prescreening forms for MR safety and gave written informed consent. The Ethical Committee of the Psychology Department at the University of Amsterdam approved the experiment. Participants were compensated 25 euros per session.

#### Experimental design

Each session consisted of 6 runs in each of which all 90 images were presented in random order in an event-related design, lasting in total 243.6 seconds each. Each scene run began with an introduction screen explaining the task to perform (locomotive action affordances, object, or fixation). Stimuli were presented at a resolution of 1017×1017 pixels such that they subtended ∼20 degrees of visual angle. Each image (independent of task) was overlaid with a fixation cross, randomly selected from one of six colors. Each image was presented for 1s followed by an ISI (randomized: 2.2, 3.8, or 5.4 seconds) with a gray screen and a white fixation cross. In each run, an additional response screen was displayed after the ISI for a subset of 15 images prompting participants to indicate their selection with a button press. These response-prompted images were pseudo-randomly-selected such that one behavioral response for each image was collected across all 6 runs. The response screen was presented for 3.8 seconds displaying the six response options specific to each task. The three tasks were adapted from the previously conducted behavioral study. In the locomotive action affordance task, participants were instructed to press the same response options as in the behavioral experiment (Walking, Biking, Driving, Climbing, Swimming, Boating) during the response trials. In the object task, participants were instructed to choose from six response options (Building, Plant, Furniture, Road, Body of Water, Stones). In the fixation task, participants were instructed to select the color (Blue, Red, Orange, Purple, Yellow, Cyan) of the fixation cross. When pressing a button, the color of the text label would turn grey indicating a pressed button. As in the previous behavioral study, participants were allowed to select multiple response options per trial in the locomotive action affordance and in the object task. While a response was only required for a subset of trials in each run, response screens appeared in random order ensuring participants needed to perform the task during each trial. The task session orders were counterbalanced across subjects, and all subjects completed all four sessions each scheduled on different days without any specific time frame between the sessions. Every experimental session took approximately 1.5 hours.

#### MRI acquisition

All participants were scanned on a Philips Achieva 3T MRI scanner and a 32-channel SENSE head coil. Each scanning session started with a survey scan for spatial planning of subsequent scans, followed by a structural T1-weighted scan acquired using a T1TFTE pulse sequence (TR = 8.262 ms, FOV = 240 X 220 X 188 mm (AP x FH x RL) with a resolution of 1 x 1 x 1 mm voxel size). During localizer scans (session 1, see ’Functional localizers’ below), functional images were acquired using a gradient echo, echo-planar pulse sequence (TR = 2,000 ms; TE = 27.63 ms; FA = 76.10; 36 sagittal slices with interleaved acquisition; 3 x 3 x 3 mm voxel size; 80 x 80 matrix; 240 x 240 x 118.5 mm FoV) covering the whole brain. A total of 140 volumes were recorded. In the same session, a set of population receptive field mapping runs were collected (not further analyzed in this study) using the same sequence, whereby a total of 150 volumes were recorded. In the following three main experimental task sessions, another T1-weighted scan was acquired in the beginning of the session followed by functional T2^∗-^weighted sequences using a multiband gradient echo EPI sequence (TR = 1,600 ms; TE = 30 ms; FA = 70; 56 sagittal slices with the interleaved acquisition; 2 x 2 x 2 mm voxel size; 112 x 112 matrix; 224 x 224 x 123 mm FoV) covering the whole brain. A total of 281 volumes were recorded per run.

#### Functional localizers

To identify category-selective regions of interest (ROIs), participants completed 6 functional mapping runs, consisting of a block design, each lasting 6.9 minutes (208 TRs) (same stimuli and paradigm as in Silson et al. (2022, 2019)). Twenty color images (568 x 568 pixels) from one of six categories were displayed in each block: Faces, Objects, Body Parts, Buildings, Scenes, and Scrambled Images. Participants performed a one-back repetition detection task while stimuli were shown against a gray background for 300 ms with gaps of 500 ms in blocks of 16 seconds duration. Blocks were separated by 4-second short fixation breaks and 16-second long fixation breaks at the start and end of the runs. Category block orders were counterbalanced within and between runs to ensure equal occurrence and spacing in a mirror-symmetrical order.

#### fMRI preprocessing

Preprocessing was performed using *fMRIPrep* 23.1.4 ((Esteban et al., 2019); (Esteban et al., 2018, RRID: SCR 016216) which is based on *Nipype* 1.8.6 ((Gorgolewski et al., 2011); (Gorgolewski et al., 2018, RRID: SCR 002502)). The 4 T1-weighted (T1w) images collected for each subject were corrected for intensity nonuniformity (INU) with N4BiasFieldCorrection (Tustison et al., 2010), distributed with ANTs (Avants et al., 2008, RRID:SCR 004757). The T1w-reference was then skull-stripped with a *Nipype* implementation of the antsBrainExtraction.sh workflow (from ANTs), using OASIS30ANTs as target template. An anatomical T1w-reference map was computed after registration of the 4 T1w images (after INU-correction) using *mri robust template* (FreeSurfer 7.3.2, Reuter et al., 2010). Volume-based spatial normalization to one standard space (MNI152NLin2009cAsym) was performed through nonlinear registration with antsRegistration (ANTs), using the brain-extracted versions of both T1w reference and the T1w template. The following template was were selected for spatial normalization and accessed with *TemplateFlow* (23.0.0, Ciric et al., 2022): *ICBM 152 Nonlinear Asymmetrical template version 2009c* (Fonov et al., 2009, RRID:SCR 008796; TemplateFlow ID: MNI152NLin2009cAsym).

#### Functional data preprocessing

For each of the functional runs per subject (across all tasks and sessions), the following preprocessing was performed. First, a reference volume and its skull-stripped version were generated using a custom methodology of *fMRIPrep*. Head-motion parameters with respect to the BOLD reference (transformation matrices, and six corresponding rotation and translation parameters) were estimated before any spatiotemporal filtering using *mcflirt* (FSL, Jenkinson et al., 2002). A B0-nonuniformity map (or fieldmap) was estimated based on one echo-planar imaging (EPI) references with *topup* ((Andersson et al., 2003); FSL None). The estimated fieldmap was then aligned with rigid-registration to the target EPI (echo-planar imaging) reference run. The field coefficients were mapped onto the reference EPI using the transform. The BOLD reference was then co-registered to the T1w reference using *bbregister* (FreeSurfer) which implements boundary-based registration (Greve and Fischl, 2009). Co-registration was configured with six degrees of freedom. Several confounding time-series were calculated based on the preprocessed BOLD: framewise displacement (FD), DVARS and three region-wise global signals. Frames that exceeded a threshold of 0.5 mm FD or 1.5 standardized DVARS were annotated as motion outliers. Gridded (volumetric) resamplings were performed using *antsApplyTransforms* (ANTs), configured with Lanczos interpolation (Lanczos, 1964). All resamplings were performed with a single interpolation step by composing all the pertinent transformations (i.e. head-motion transform matrices, susceptibility distortion correction when available, and co-registrations to anatomical and output spaces). Many internal operations of *fMRIPrep* use *Nilearn* 0.10.1 (Abraham et al., 2014a, RRID:SCR 001362), mostly within the functional processing workflow. For more details of the pipeline, see https://fmriprep.readthedocs.io/en/latest/workflows.html.

#### Localizer fMRI analysis and ROI definitions

General linear models (GLM) were created for each individual subject and each individual run using the implementation provided by *Nilearn/Nistats* (https://nilearn.github.io; Abraham et al., 2014b, version 0.9.0). This GLM included 16s block regressors for each of the six categories (scenes, buildings, objects, faces, bodies, scrambled) which were convolved with a canonical hemodynamic response function (Glover, 1999) and fitted using AR1 autocorrelation correction and smoothed with a Gaussian 3mm FWHM filter. Subject-specific scene-selective regions were identified by first generating thresholded (p *<*0.05, FDR) z-scored t-statistic activation maps to a number of contrasts that tested for higher activity to scenes than to objects, faces and scrambled image blocks (e.g. ’scenes’ *>*’faces’ & ’objects’ and ’scenes’ *>*’scrambled’) which were subsequently intersected with an atlas of scene-selective parcels from Julian et al. (2012). In the last step, all contrast activation maps were unified to create subject-specific ROIs for PPA, OPA, MPA. The obtained ROIs contained on average 71% (PPA), 78% (OPA) and 66% (MPA) of the voxels in the atlas parcels, resulting in an average of 915 (PPA), 302 (OPA) and 1868 (MPA) voxel-sized ROIs.

#### Experimental task sessions fMRI data analysis

The fMRI time-series data from the main experimental sessions were fitted with subject- and run-specific GLMs which contained one regressor for each of the 90 images convolved with a canonical hemodynamic response function (Glover, 1999). Given the randomization of response prompts across runs and the temporal separation of scene stimulus and response prompt screen, images with and without response prompts were not modeled separately. Additionally, 3 translational motion regressors, 3 rotational motion regressors and 3 low-frequency cosine DCT-basis regressors calculated by fMRIPrep were included as covariates. GLMs were fitted using an AR1 autocorrelation correction and smoothed with a Gaussian 3mm FWHM filter, after which one contrast per image was computed and resulting beta-estimates were masked by the subject-specific ROIs. The resulting voxel patterns were averaged across runs and used to compute subject-specific RDMs for each ROI using 1-Pearson correlation distances between each pair of images.

#### In-scanner behavioral data analysis

For each participant, individual RDMs were calculated using 1-Pearson correlation distance and standardized features consisting of the annotations made by each participant indicating the possibility for the six locomotive actions, the presence of objects, or the color of the fixation cross.

### DNN feature activations

To derive RDMs based on deep neural network features, we extracted activations from a number of pretrained DNN models for our full image set using the Net2brain Python package (https://github.com/ cvai-roig-lab/Net2Brain; Bersch et al., 2022, version 0.1.0). We selected 13 commonly used DNNs (see **Supplementary Table 1** for an overview), divided in seven different groups (see legend of **Figure 5A**). We extracted feature activations for the layers pre-selected by Net2Brain and computed RDMs of the standardized features per layer (removing the mean and scaling to unit variance) using 1-Pearson correlation distance metric.

### End-to-end training and fine-tuning on affordances

Given the limited alignment of pre-trained DNNs with action affordances, we also attempted to directly train models on scene affordances and fine-tune pre-trained models specifically for this task. We used the functions subset of the SUN Attribute Dataset (Patterson et al., 2014), which provides labels for different scene attributes. Each label is annotated with a number of votes ranging from zero to three: a score of zero indicates the absence of an affordance, while a score of three signals its guaranteed presence. Here, an affordance was considered present in an image if it had received at least two votes by the annotators, and absent if it had received zero votes. Affordances that received a single vote were excluded due to their ambiguity. We also omitted affordances with fewer than 300 instances from our training set, as this was considered insufficient for training deep convolutional neural networks effectively. This process resulted in a dataset comprising of 8,147 images and 24 of 37 affordance labels (see **Supplementary Table 4 for overview**. For the models, we utilized a ResNet50 and the Vision Transformer (ViT) Base Patch 16, both implemented in PyTorch 1.11.0. The ResNet50 was imported from the Net2Brain Toolbox (Bersch et al., 2022) and the ViT Base Patch 16 from the ”PyTorch Image Models” repository by (Steiner et al., 2021). We adjusted both models to feature a final layer with 24 output units, each representing an affordance label, and included a sigmoid activation function and an Adam optimizer. We employed Binary Cross Entropy (BCE) Loss since each label was treated as an independent binary classification task (presence vs absence). While for end-to-end training the models learned entirely from the training data, for the fine-tuning approach we loaded pre-trained weights and only updated the final fully connected layer while keeping all other layers frozen. We applied 5-fold cross-validation to ensure the robustness of our optimization and utilized early stopping based on validation loss to prevent overfitting. Our training protocol used a batch size of 64, and we optimized the models using a two-phase hyperparameter tuning process: we first employed a random search to quickly identify effective parameter ranges, followed by a grid search to fine-tune these parameters and select the best-performing models. **Supplementary Table 6** shows some of the parameter configurations of the trained and fine-tuned models.

### GPT-4 annotations

We presented each of our 231 images to GPT-4 OpenAI et al. (2023) to generate various sentence captions and behavioral responses similar to those generated by the human participants for locomotive action affordances and objects (see **Supplementary Table 2** and **Supplementary Table 3** for an overview of used prompts.). For the sentence captions, we prompted for unique captions for our representational spaces focusing on action affordances and objects and general captions as a control. For the behavioral responses, the tasks were to pick a single affordance/object, multiple affordances/objects and create a bag of words representation of all possible affordances and objects in the image. For the sentence caption RDMs each caption was passed through an embedding model (all-MiniLM-L6-v2, (Reimers and Gurevych, 2019)) and the resulting feature vectors were standardized (removing the mean and scaling to unit variance) and RDMs created using 1-Pearson correlation distance metric. The behavioral response RDMs were created based on a one-hot encoding of the resulting responses using Euclidean distance metric for the single annotation behavior and 1-Pearson correlation distance for the multiple and BoW annotations.

### Environment Model RDMs and GIST model

Computer vision datasets often put images in three different types of environments. Indoor, outdoor manmade and outdoor natural. We used these types of environments for each image to create three different binary model RDMs. One representing Indoor *vs.* Outdoor scenes (Outdoor manmade, Outdoor natural). One natural *vs.* manmade (Indoor, Outdoor manmade) and one with all three types of environments separate. RDMs were created based on a one-hot encoding of the environment types and using the Euclidean distance metric. Prior research has shown that a spatial envelope (GIST) (Oliva, 2005) accounts for significant variance in fMRI and MEG responses across the visual cortex Ramkumar et al. (2016); Watson et al. (2014, 2017) we also included a model RDM to control for these. We used a 194×194 window grid across each image to capture dominant orientation contrasts at three different spatial frequencies. The number of orientations changes with spatial frequency, with 8 orientations at the highest frequency, 6 at the mid-range, and 4 at the lowest. Resulting in 512-dimensional feature vectors. These were used to create the GIST RDM using 1-Pearson correlation distance.

### Representational Similarity Analysis

RDMs were compared to one another by computing Spearman’s *ρ* as proposed by the RSA toolbox (Kriegeskorte, 2008; Nili et al., 2014) (lower triangle, excluding the diagonal; Ritchie et al. (2017)). This was done on the subject average in-scanner behavior and single subject ROI RDMs. Noise ceilings for fMRI RDMs and RDMs derived from behavioral annotations were computed following the approach recommended by Nili et al. (2014). The upper bound of the noise ceiling was computed by calculating for each participant the Spearman correlation between their own, individual RDM and the group-average RDM across all participants, and averaging the resulting correlations. The lower bound was computed using a leave-one-out approach, correlating each participant’s RDM with the average RDM of all other participants, and averaging the resulting correlations. We compute layer-wise comparisons of DNN feature activation RDMs with single-subject ROI RDMs and single-subject in-scanner behavior RDMs always reporting the highest correlating layer (see **Supplementary Table 5** and **Supplementary Table 1** for all extracted layer)s. One-sample t-tests were used to test whether the average correlation was significantly different from zero, and paired sample t-tests were used to test for significant differences between correlations. The obtained p-values were corrected for multiple comparisons using Bonferroni correction (2 for fMRI: 0.05/2 = 0.025, 3 for fMRI task-effect: 0.05/3 = 0.017, 13 DNN models: 0.05/13 = 0.003). Pairwise comparisons were also corrected for multiple comparisons using Bonferroni correction when comparing different tasks and DNN models to one another. When comparing DNN features with behavioral annotation RDMs obtained from the online behavioral experiment and the automatic labeling, we used (n = 1000) bootstrapping (sampling rows/columns with replacement) to obtain the 95 % confidence intervals. The differences between the bootstrapped distributions were statistically compared using two-tailed sign test and FDR-corrected for multiple comparisons.

#### Whole-brain searchlight analysis

To investigate affordance processing beyond scene-selective ROIs, we performed a series of whole-brain searchlight analysis. RDMs were computed using correlation distance metric based on multivoxel patterns within 5-mm radius spherical searchlights extracted, using the Nilearn (Abraham et al., 2014b) package. Consistent with the ROI analyses, we calculated partial correlations for locomotive affordances, controlling for contributions from the GIST model and object annotations. Spearman correlation values from the RSA were assigned to the center voxel of each searchlight, generating whole-brain maps of locomotive affordance coding. Individual subject maps were smoothed with an isotropic Gaussian kernel (8-mm FWHM). Group-level statistical significance was evaluated using one-sample t-tests, with p-values thresholded through permutation testing (n = 10,000) at a negative log10 p-value threshold of 1.3, equivalent to p *<*0.05. (Smith and Nichols, 2009)

### Hierarchical clustering and Variance Partitioning

To visualize different aspects of the representational structure of the RDMs, we applied hierarchical clustering using clustermap in seaborn (Waskom, 2021, version 0.12.2) which uses the hierarchical/agglomerative clustering algorithm implemented in scipy (Virtanen et al., 2020, version 1.8.0). The default settings were used, applying a hierarchical clustering using ”average” which computes clusters using a nearest-neighbor chain algorithm with the Euclidean distance metric until all data points are clustered. For environment-based sorting, we used the image names categorized as indoor, outdoor natural, and outdoor man-made. The same clustering method was also used for the sorting by locomotive action affordances. To explore if locomotive action affordances can be explained by other visual features like material and object annotations, we performed variance partitioning based on multiple linear regression. For this analysis, we used the off-diagonal of the locomotive action affordance annotations from the online experiment as the dependent variable and the behavioral ratings from the other scene properties (objects, materials, and global properties) as independent variables. We selected these three features as a model with these three variables reached the highest explained variance with the locomotive action affordance annotations. To obtain unique and shared variance across the variables multiple regression analyses with different feature compositions are computed. A full regression model with all features, individual models with single features, and all feature pairs. From these unique and shared variance was estimated. For this we used the vegan package (Oksanen et al., 2001, version 2.6-6.1) in R (R Core Team, 2021, version 3.6.1). For visualization purposes, we created an Euler diagram, using EulerAPE (Micallef and Rodgers, 2014).

## Code availability

Stimuli, behavioral annotations, MRI-data and extracted DNN feature activations will be made available on the OSF https://osf.io/v3rcq/. Python scripts containing preprocessing and analysis code to reproduce the results will be available at https://github.com/cgbartnik.

## Acknowledgements

This work was supported by an VENI grant (VI.Veni.194030) from the Netherlands Organisation for Scientific Research (NWO) to IIAG. We thank Lukas Muttenthaler, Steven Scholte and Chris Baker for helpful feedback.

## Supplementary Information

**Supplementary Figure 1:**
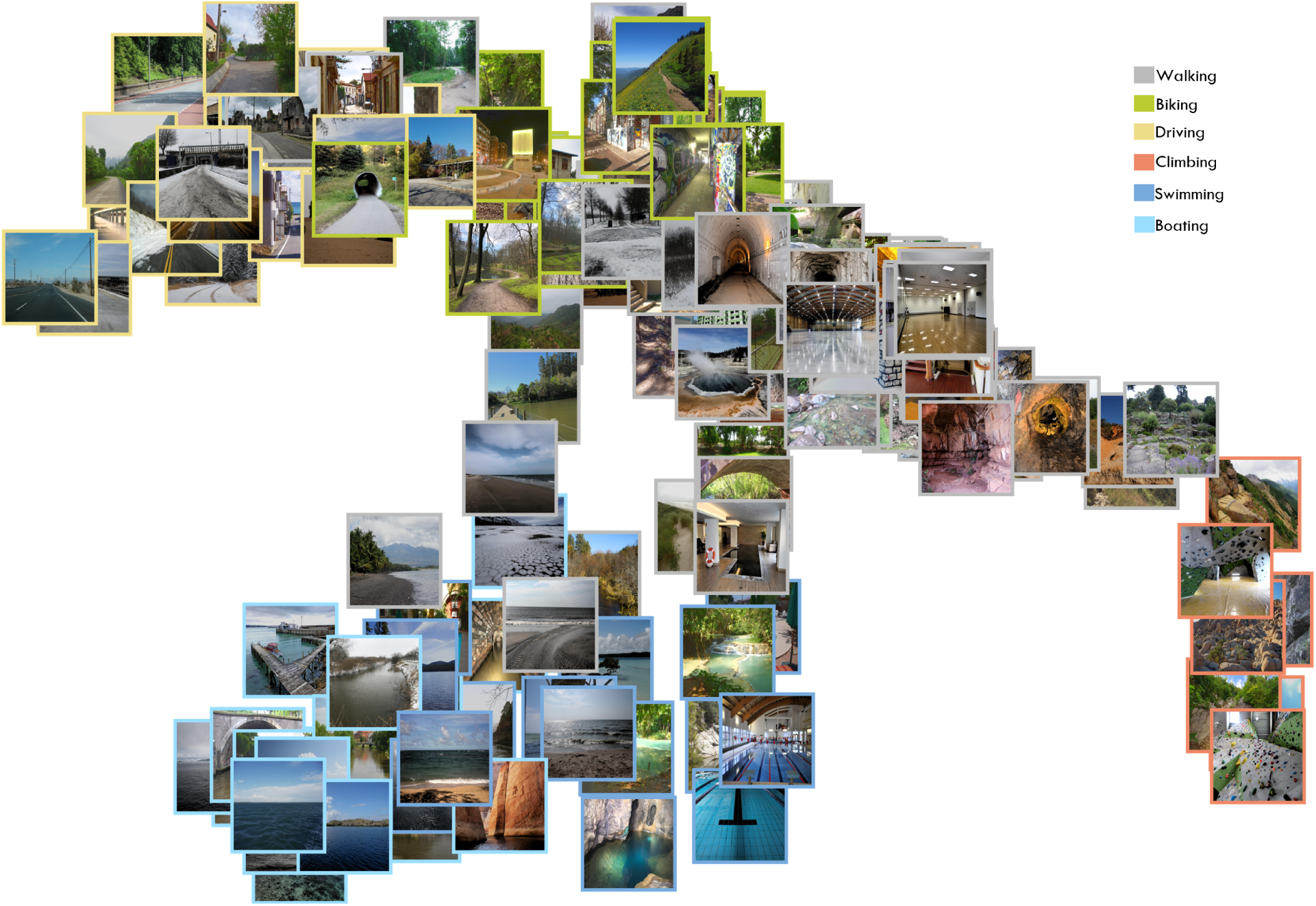
Two-dimensional multidimensional scaling of locomotive action affordance representational space. This figure displays the same spatial arrangement of our complete 231 stimulus set as displayed in Figure 2 but with stimulus thumbnails each framed by a colored border. The color coding corresponds to the most frequently chosen locomotive action associated with each image (see Materials and Methods for details).

**Supplementary Figure 2:**
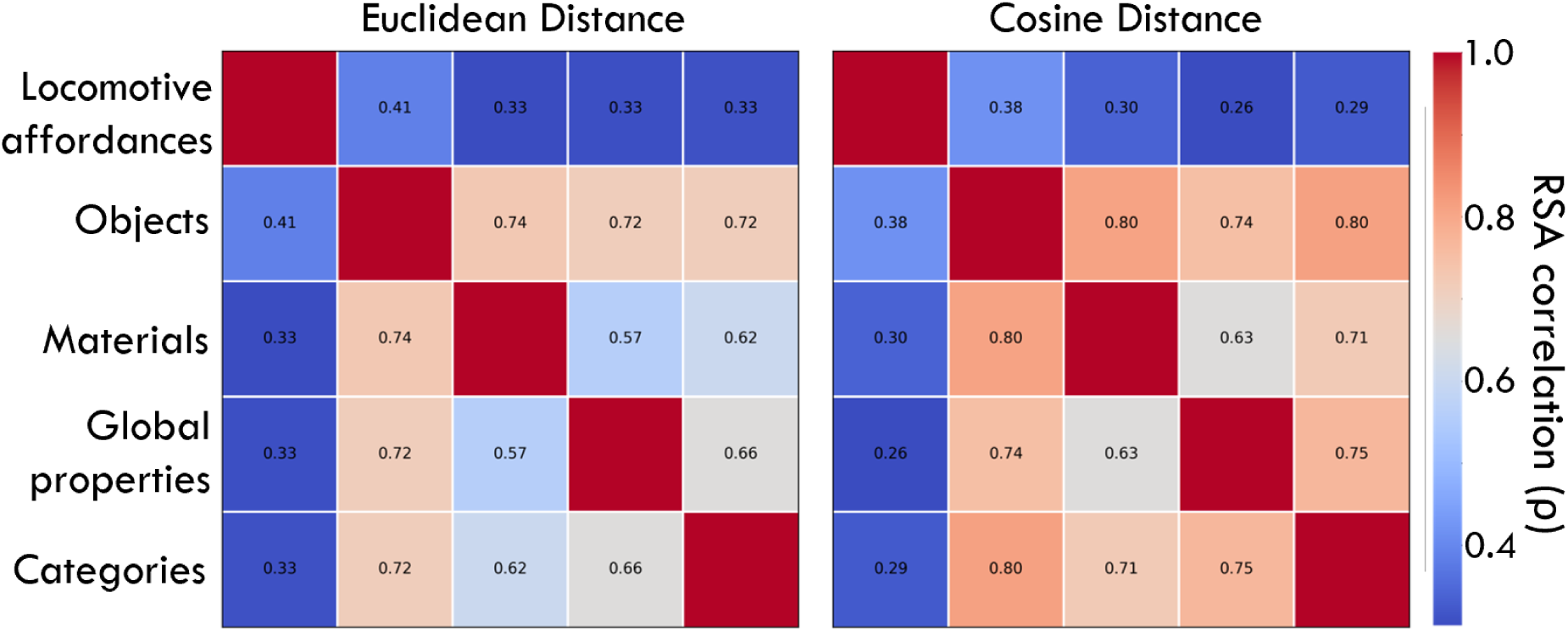
Correlation between locomotive action affordances and other scene properties using different distance measures to compute RDMs. We consistently find that locomotive action affordances are a unique space and that the other visual scene properties highly correlate with each other, also when behavioral RDMs are constructed using Euclidean or cosine distance.

**Supplementary Figure 3:**
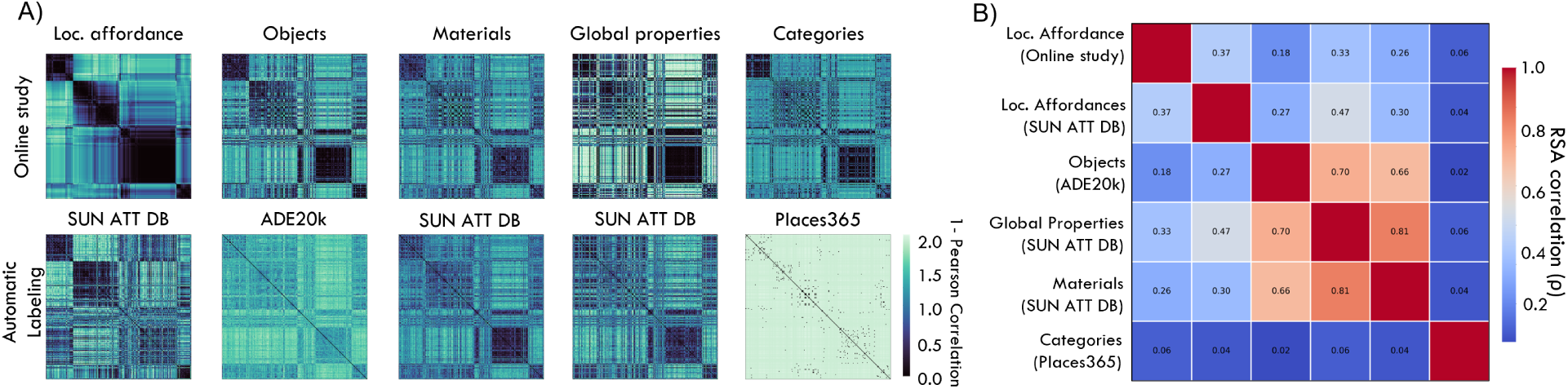
Comparison of RDMs for all scene properties obtained using behavioral annotations or automated labeling methods. **(A)** Upper row, RDMs of the representational space of various scene properties for which we collected annotations in an online experiment. Bottom row, RDMs of similar spaces that were created using automatic labeling by utilizing largescale image datasets and computational models. These were used to evaluate if the interrelation of the RDMs that we found in our behavioral experiment were a product of our limited selection of possible labeling options or if this is also true in more unrestricted cases. **(B)** Correlations between RDMs for all spaces computed from automatic labels, and our behavioral annotations of locomotive action affordances. We find a similar pattern as for annotations obtained through human behavior: locomotive action affordances correlate relatively low with other scene properties which highly correlate with each other. Note that category labels here are an exception, correlating poorly with all other spaces (including locomotive action affordances), unlike in our behavioral experiment. This is likely due to the high sparsity of the Place365-derived category RDM.

**Supplementary Figure 4:**
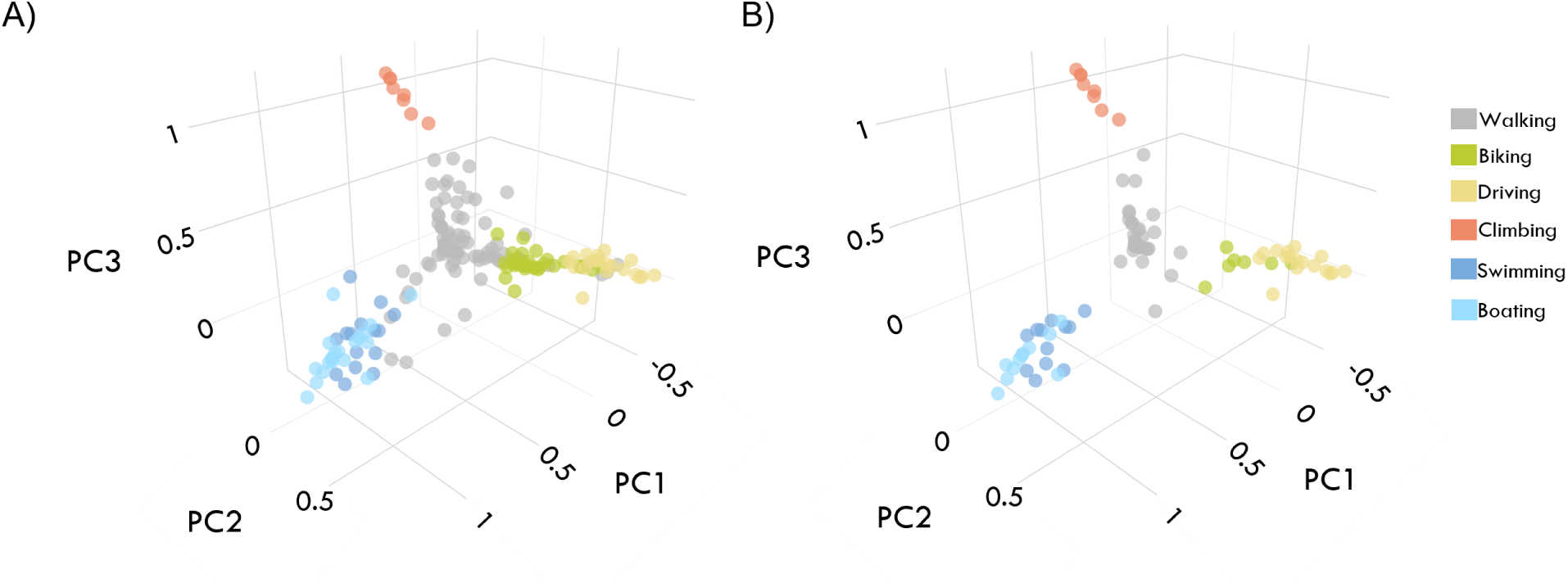
Using Principal Component Analysis for strategically sampling a stimulus subset. **(A)** PCA of the behavioral locomotive action affordance RDM from the online study including all 231 images. The color coding corresponds to the most frequently chosen locomotive action associated with each image (see Materials and Methods for details) **(B)** Highlighting the strategically chosen subset of 90 images employed in the fMRI study in the same PCA space. This subset is composed of an equal number of indoor, outdoor-natural, and outdoor-manmade scenes (30 each), selected to capture the three-dimensional locomotive action affordance structure.

**Supplementary Figure 5:**
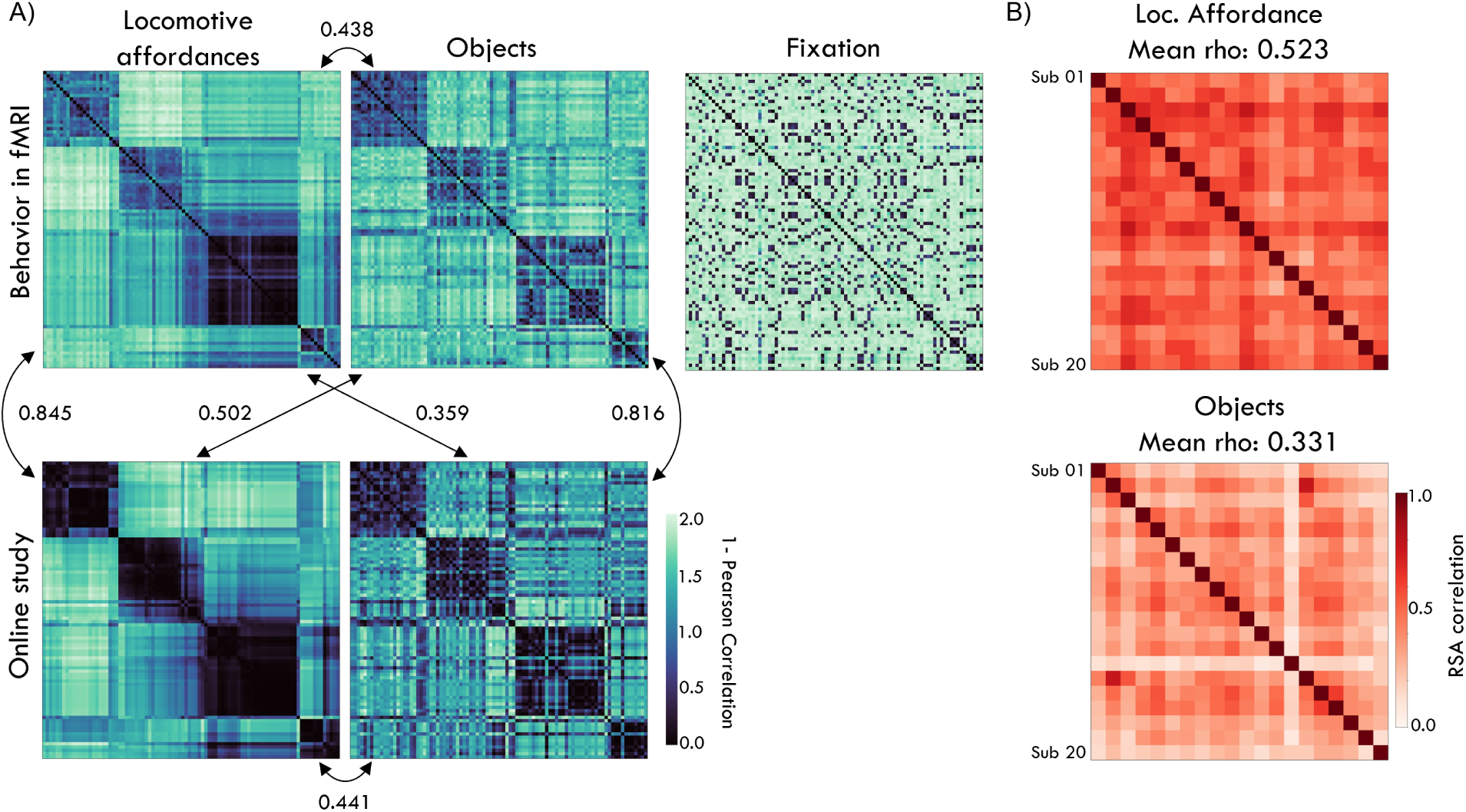
Comparison of behavioral data obtained in the online study versus fMRI experiment. **(A)** The upper row contains the average RDMs computed based on three behavioral annotation tasks (locomotive action affordances, objects and color of fixation cross) performed by the participants in the fMRI scanner. The bottom row shows the corresponding RDMs from the online study. These results show that the task replicates well in the scanner, showing a high correlation between behavioral annotations from the same task in the scanner and in the online experiment, while the correlations between tasks are similar between online and fMRI scanner participants. Behavior for the orthogonal fixation task (top right) is completely unrelated to behavioral annotations of either locomotive affordance or objects, as intended. **(B)** Intercorrelation of RDMs for each behavioral tasks across the fMRI participants. Participant performance was more consistent during the locomotive affordance task than during the object task.

**Supplementary Figure 6:**
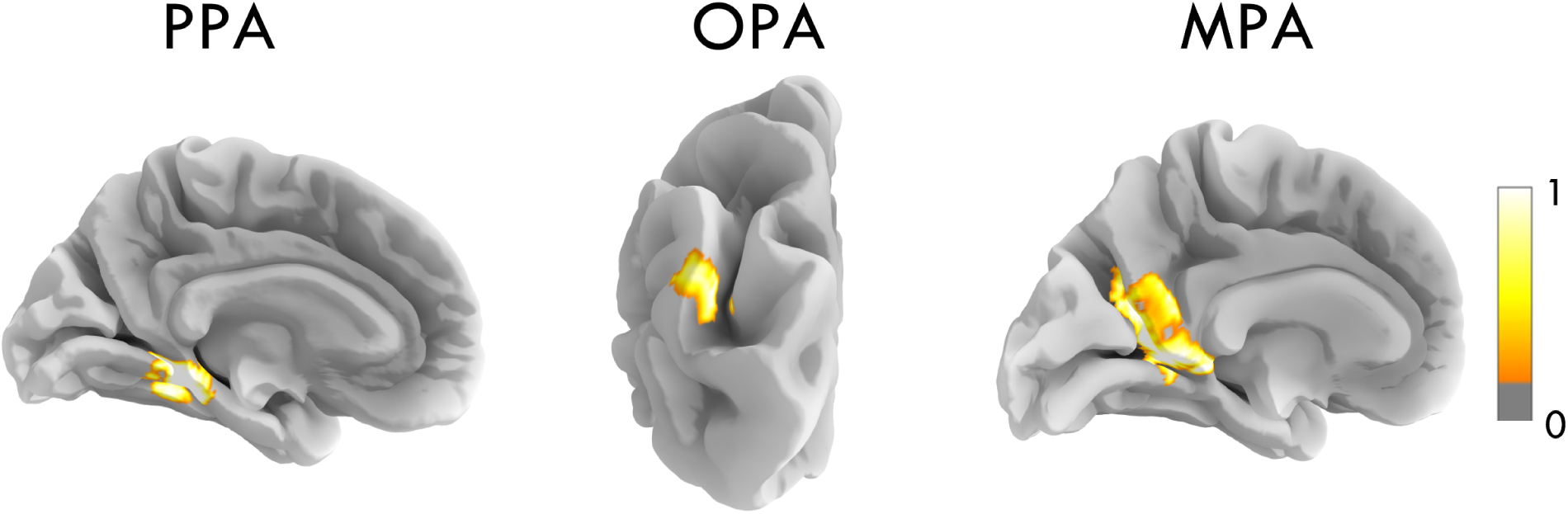
Scene-selective ROIs. Scene-selective ROI (Parahippocampal Place Area, PPA; Occipital Place Area, OPA; and Retrosplenial Cortex, MPA) masks of subject 004 in MNI-152 template space (see Materials and Methods for details).

**Supplementary Figure 7:**
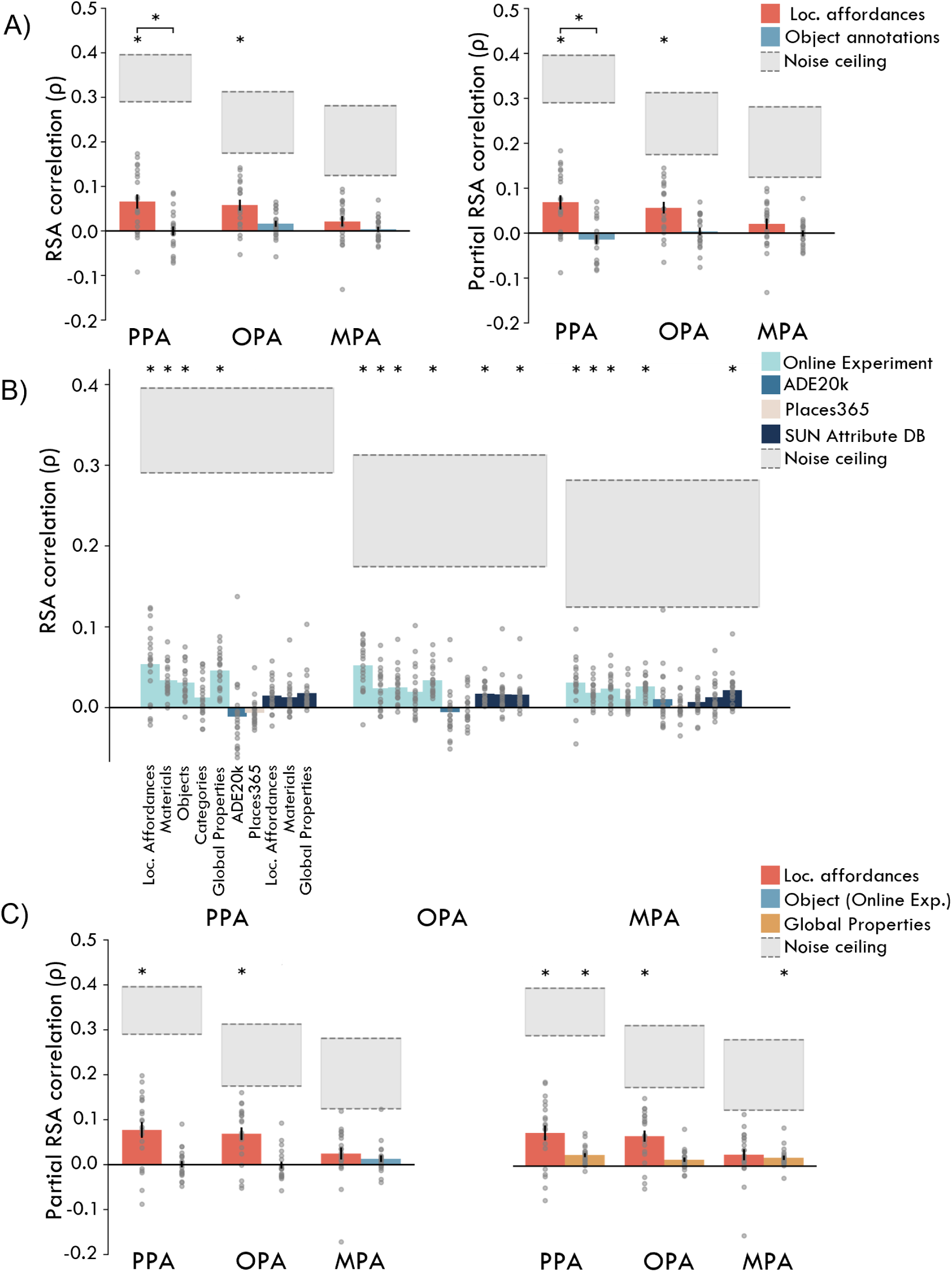
Locomotive action affordance representations in PPA and OPA for individual participants, online behavior and automatic labels. **(A)** Full (left) and partial (right) correlations of the fMRI RDMs with in-scanner individual subject behavioral RDMs. **(B)** Correlations of the fMRI RDMs with the online behavioral and the automatic labeled spaces. **(C)** Partial correlations analysis with online behavioral RDMs show unique representation of locomotive action affordances versus objects (left) and global properties (right). Error bars indicate the standard error of the mean (SEM) across participants. Asterisks (*), indicating one-sample t-test against zero p*<*0.05 corrected for multiple comparisons using Bonferroni correction.

**Supplementary Figure 8:**
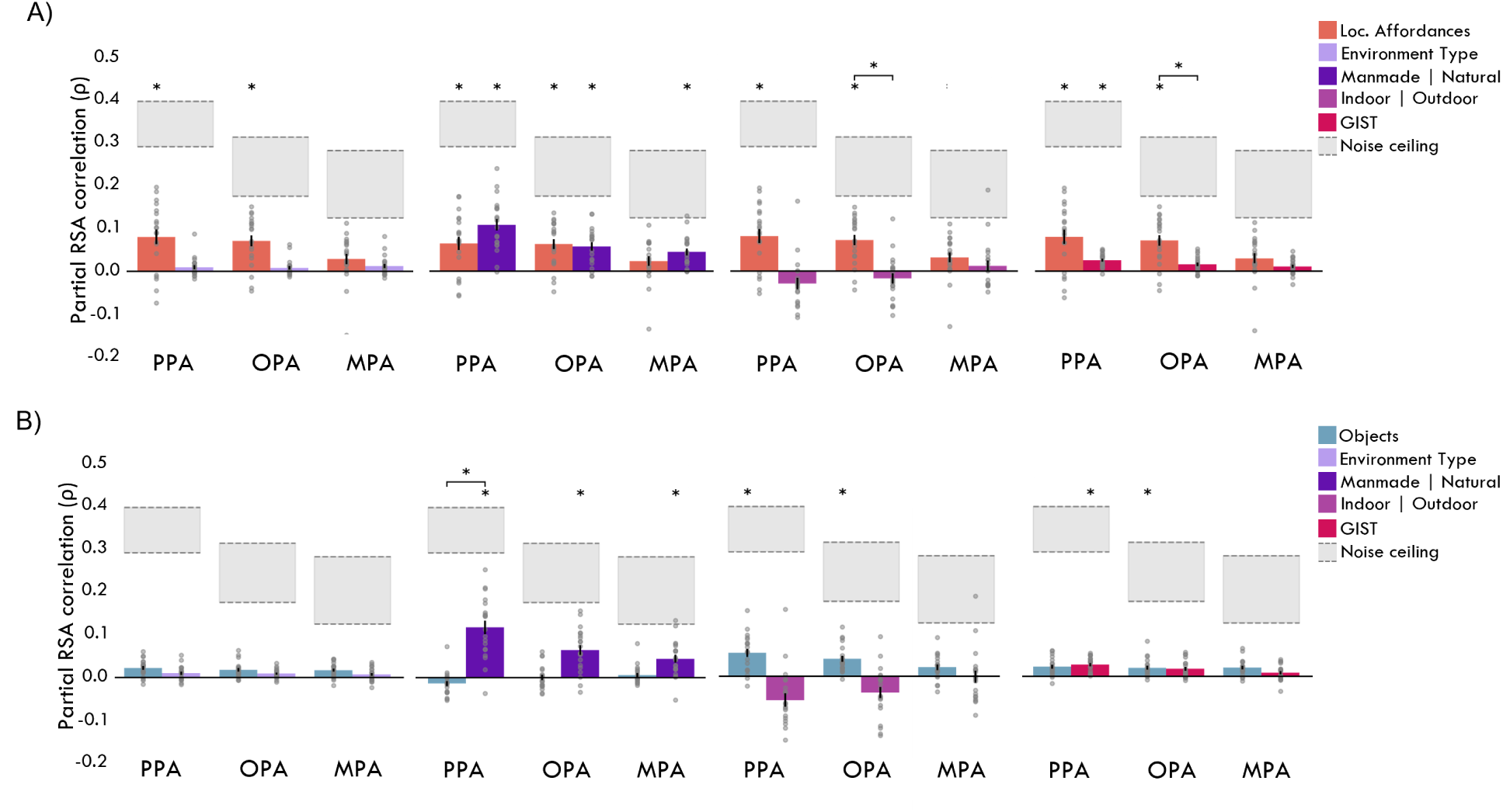
Scene-selective regions sensitive to manmade/natural environments but these can not explain locomotive action affordances. **(A)** Partial correlations of model RDMs for different environment types with locomotive action affordances behavior. First panel shows the partial correlations with a binary model RDM across all three types of environments (Indoor, Outdoor manmade, and Outdoor natural). The second panel shows the correlations with a model RDM grouping manmade (Indoor and Outdoor manmade) and Outdoor natural images. The third panel splits in Indoor and Outdoor images. The last panel shows the correlations with a model RDM derived from the GIST model using a 1024 by 1024 pixel image resolution. **(B)** Partial correlations of model RDMs for different environment types with object annotations. The same model RDMs in each panel as in (A).

**Supplementary Figure 9:**
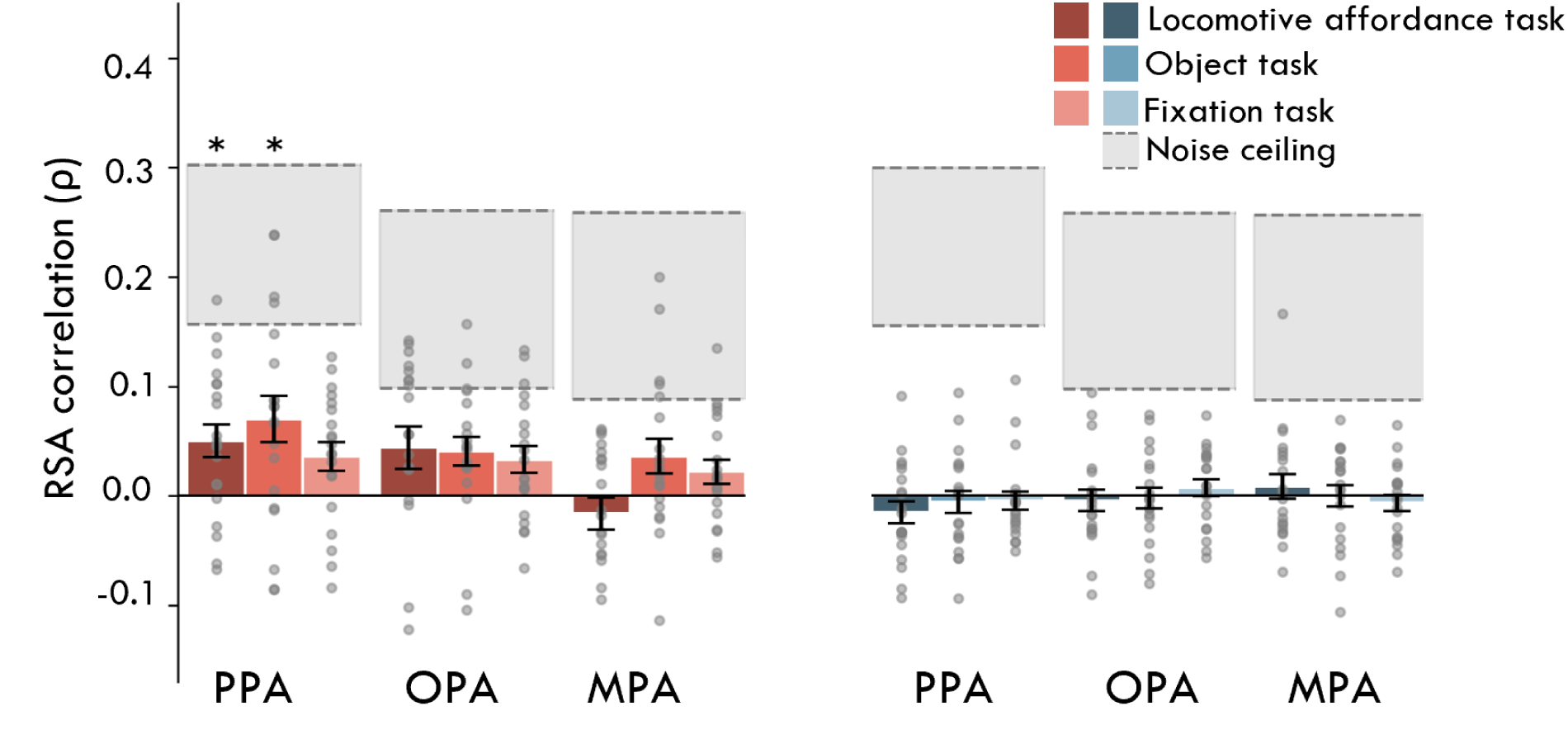
Task-specific partial correlations of fMRI RDMs with behavioral RDMs of affordance and object annotations. Partial correlations of RDMs derived from fMRI response patterns in scene-selective brain regions with locomotive action affordances (red) and object annotations (blue) RDMs during a locomotive action affordance labeling task (darker colors), and object labeling task (middle colors) and an orthogonal fixation task (light colors). Bars represent averages for individual participants (gray dots). Shaded areas delineate upper and lower noise ceilings, reflecting the range of RDM similarity between individual participants and the group mean. Error bars indicate the standard error of the mean (SEM) across participants. Asterisks (*), indicating one-sample t-test against zero p*<*0.05 corrected for multiple comparisons using Bonferroni correction.

**Supplementary Figure 10:**
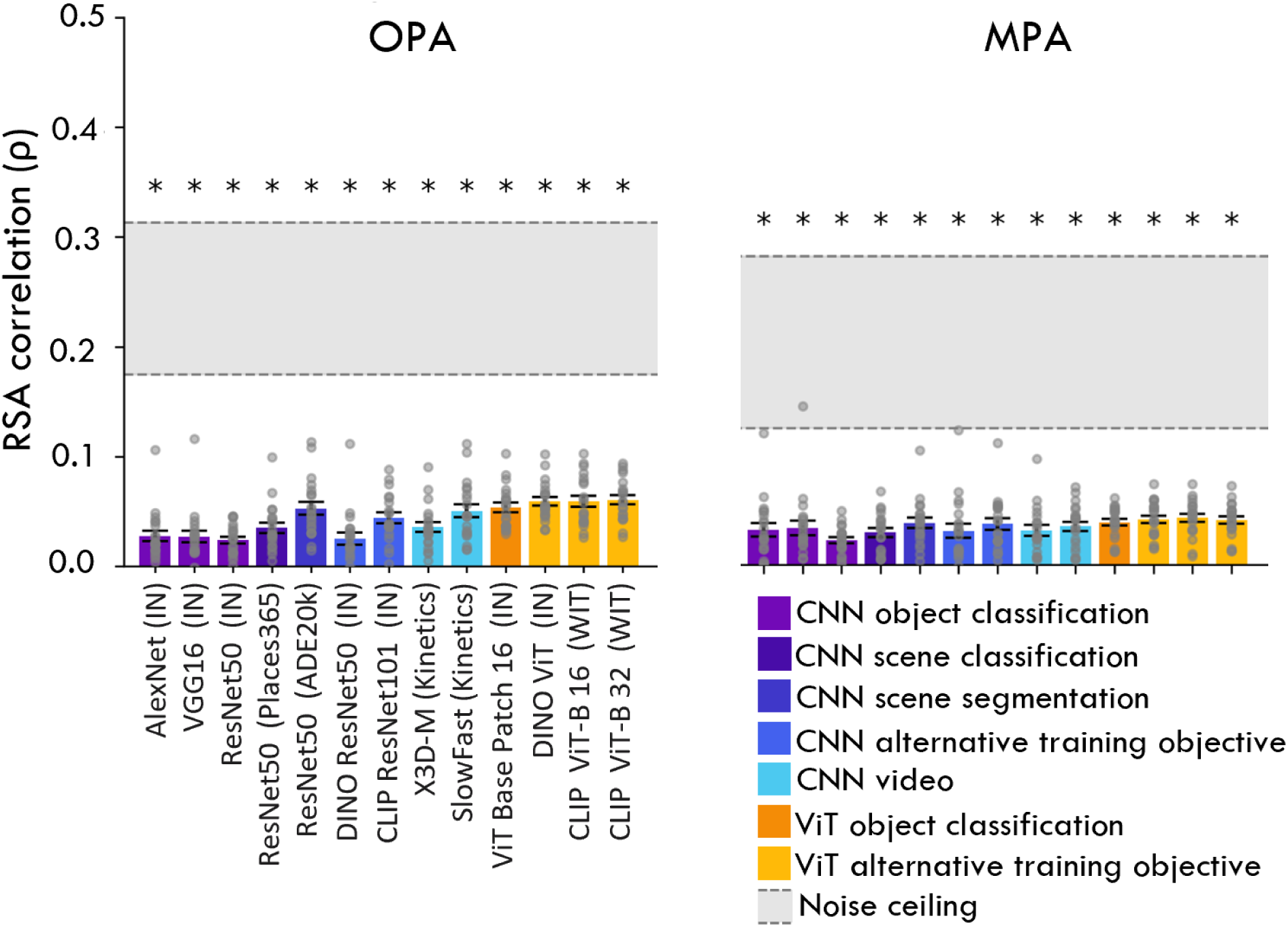
Average correlation of each DNN’s best correlating layer with representational similarity in OPA and MPA. Bars represent average correlations across participants and tasks for the best correlating layer in each model and gray dots indicate individual participants. Significant t-tests of the average correlation against zero are marked by asterisks (*), indicating p*<*0.05 corrected for multiple comparisons using Bonferroni correction. The shaded area delineates the upper and lower noise ceiling and error bars indicate the standard error of the mean (SEM) across participants.

**Supplementary Figure 11:**
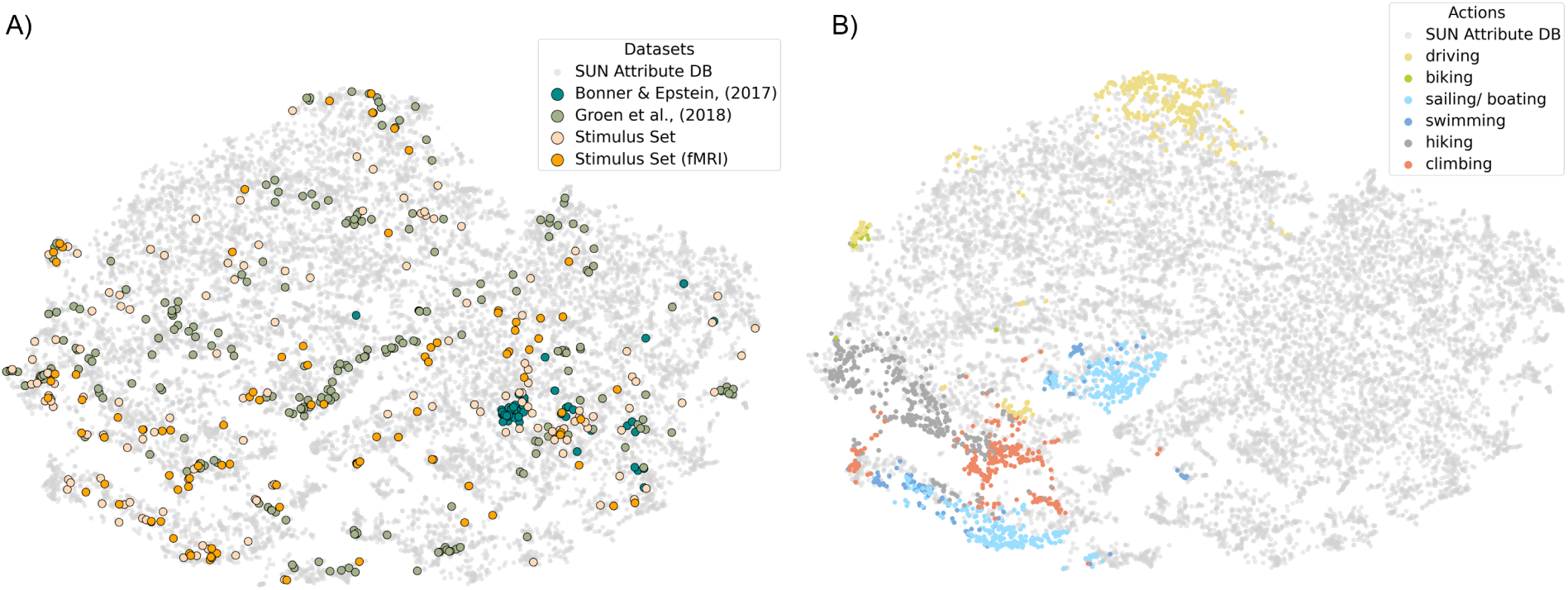
Visualization of scene representational space and associated locomotive action affordances. **(A)** t-SNE visualization of the SUN Attribute Database representational space containing over 12.000 images with various datasets from prior research (Bonner and Epstein, 2017) and (Groen et al., 2018) and our image set (whole and fMRI subset) embedded in the representational space. **(B)** t-SNE visualization of the SUN Attribute Database with coloring based if a scene was labeled to afford a specific locomotive action. These judgments are based of the labeling in the SUN Attribute Database.

**Supplementary Figure 12:**
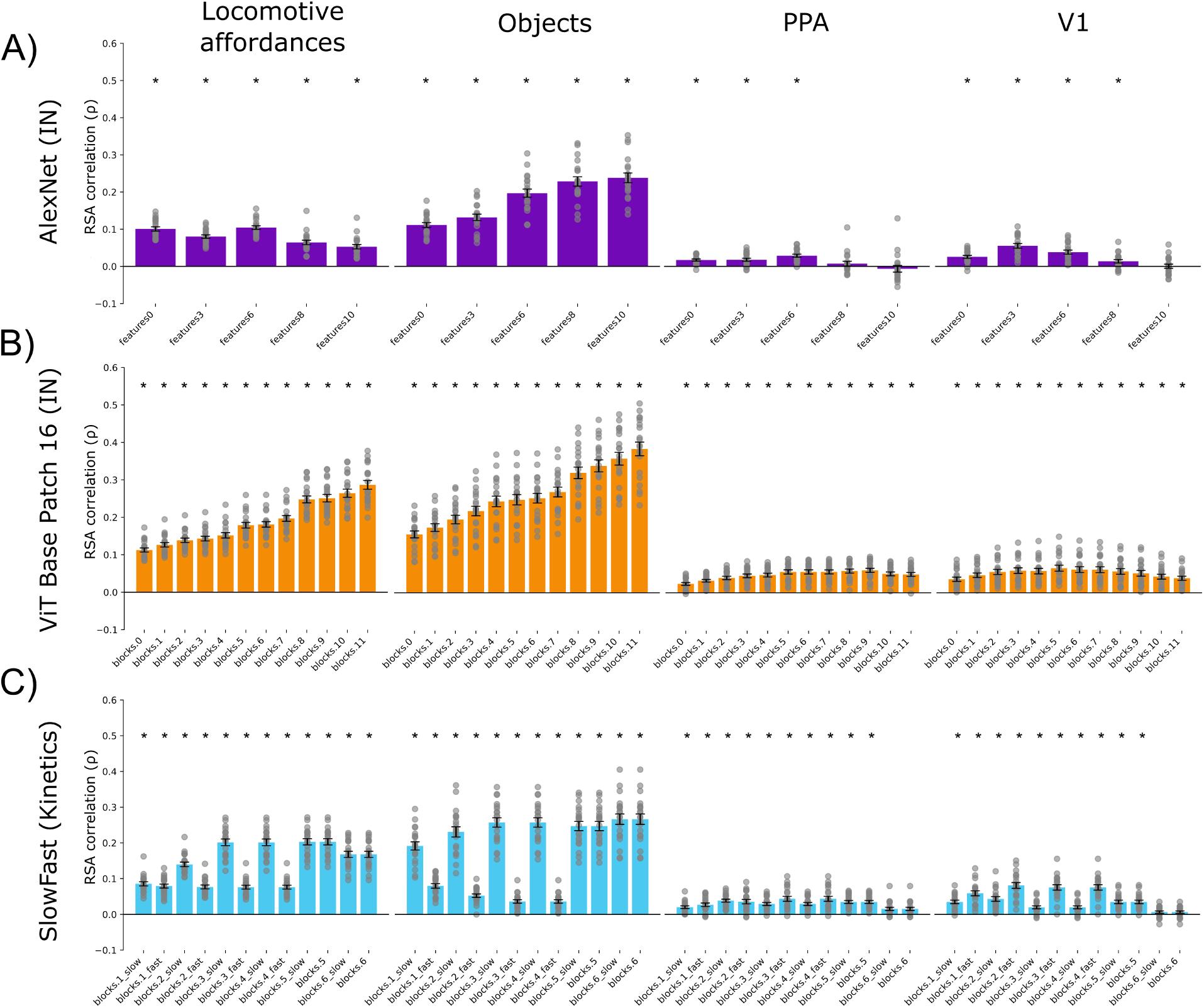
Layer-wise correlations with behavioral annotations and visual cortex ROIs. **(A)** Bars represent average correlations across of AlexNet (ImageNet) layer activations with behavioral annotations and visual cortex ROIs with error bars show the standard error of the mean (SEM) across participants. Gray dots indicate individual participants behavior and ROIs correlations. Significant t-tests of the average correlation against zero are marked by asterisks (*), indicating p*<*0.05 corrected for multiple comparisons using Bonferroni correction. **(B)** Average correlations by layer for ViT Base Patch 16 (ImageNet). Same plot elements as (A) **(C)** Average correlations by layer for SlowFast (Kinectics). Same plot elements as (A)

**Supplementary Figure 13:**
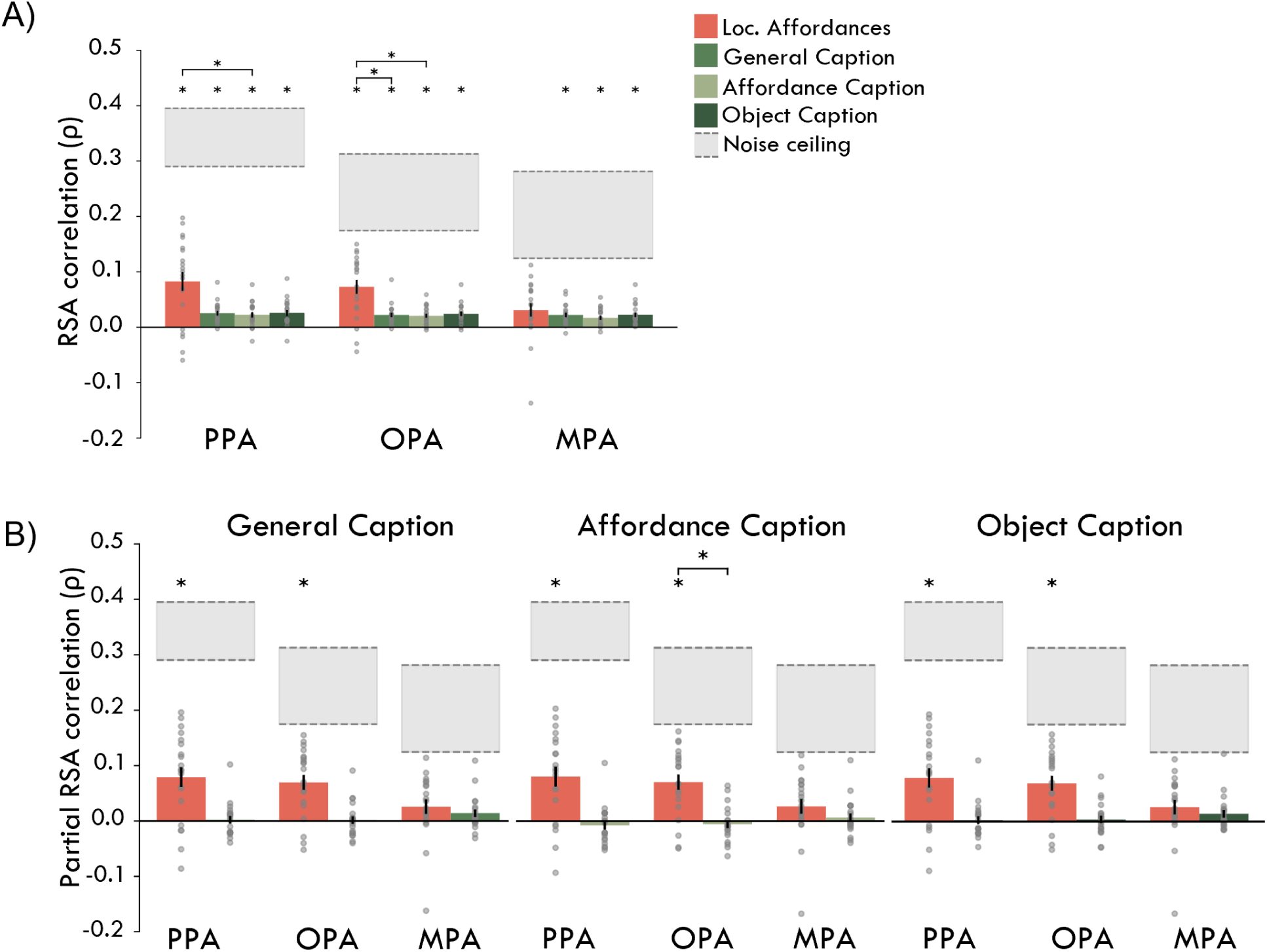
LLM-based (GPT-4) generated captions do not capture fMRI responses in scene-selective regions well. **(A)** Correlation of response patterns in scene-selective brain regions with locomotive action affordance (red) and three different GPT-4 generated captions (General Caption, Affordance Caption, Object Caption) that were transformed using the MiniLM embedding space to create representational spaces (see in **Supplementary Table 2** for details). Bars represent averages for individual participants (gray dots). Shaded areas delineate upper and lower noise ceilings, reflecting RDM similarity between individual participants and the group mean. Error bars indicate standard error of the mean (SEM) across participants. Asterisks indicate one-sample t-tests against zero (p*<*0.05) corrected for multiple comparisons using Bonferroni correction (here, 12 comparisons). **(B)** Partial correlations of the GPT-4 generated captions representational spaces with locomotive action affordances behavior. Same plot elements as (A)

**Supplementary Table 1:**
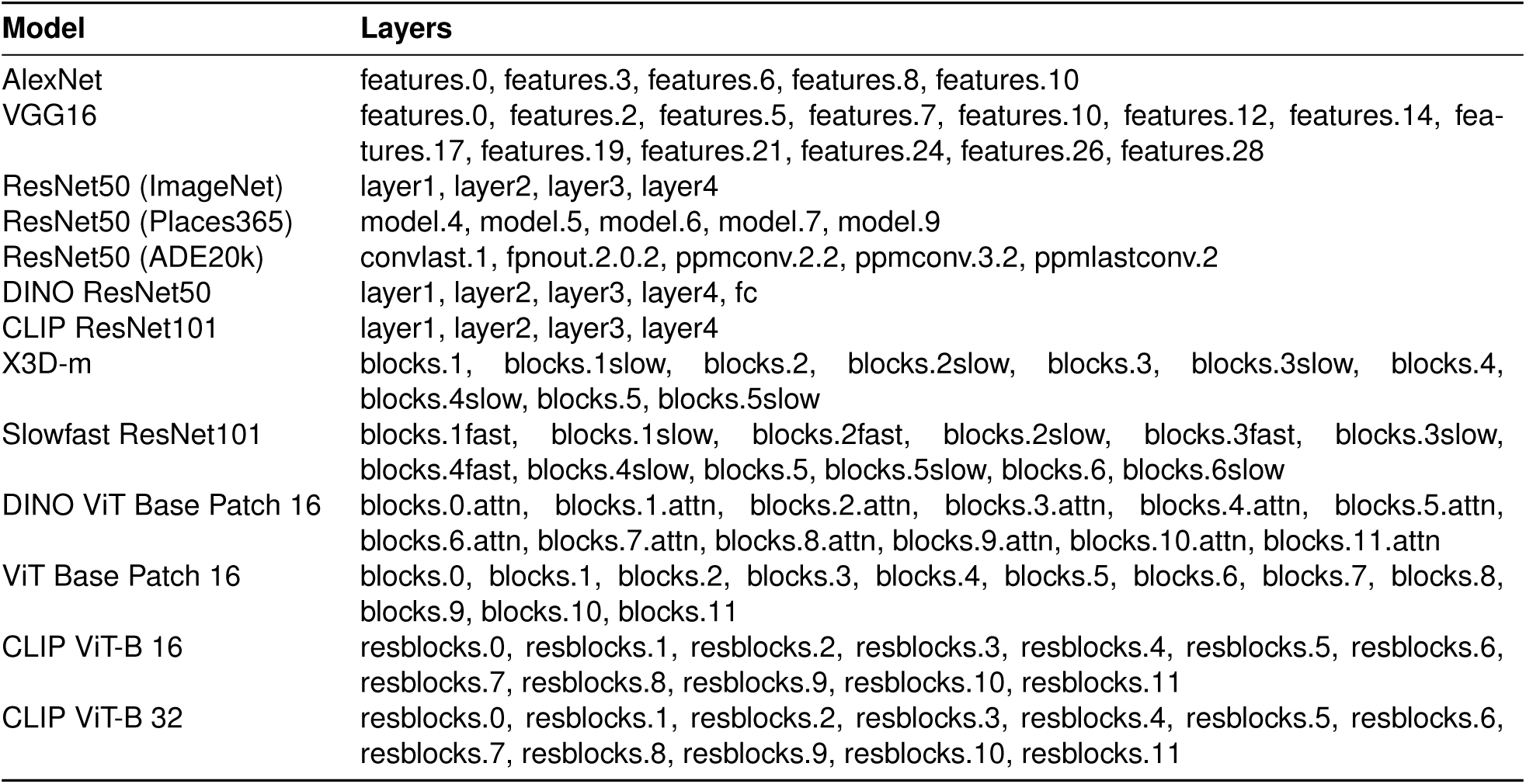
Deep Neural Network Models and the extracted layers.

**Supplementary Table 2:**
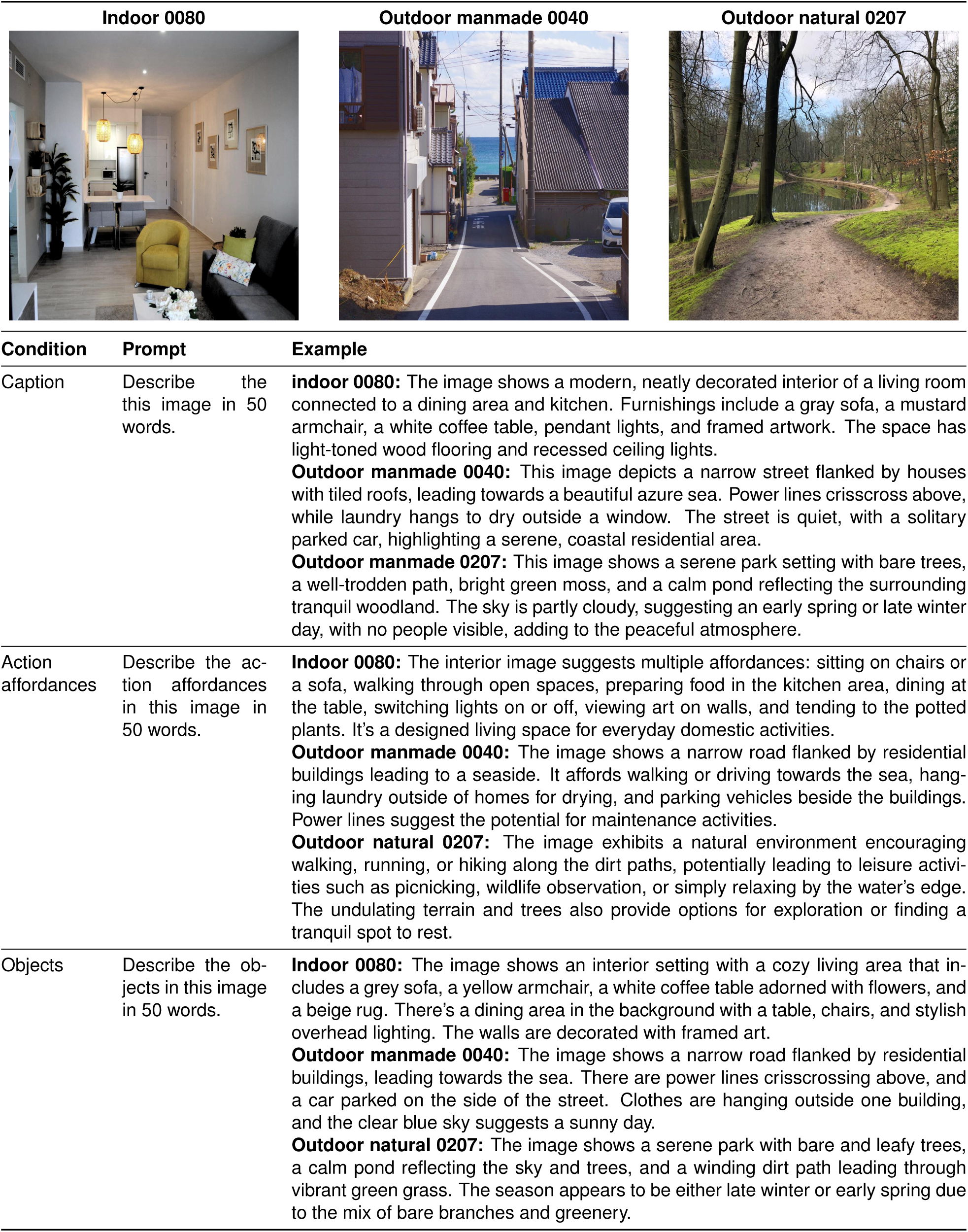
Examples of various captions generated using GPT-4.

**Supplementary Table 3:**
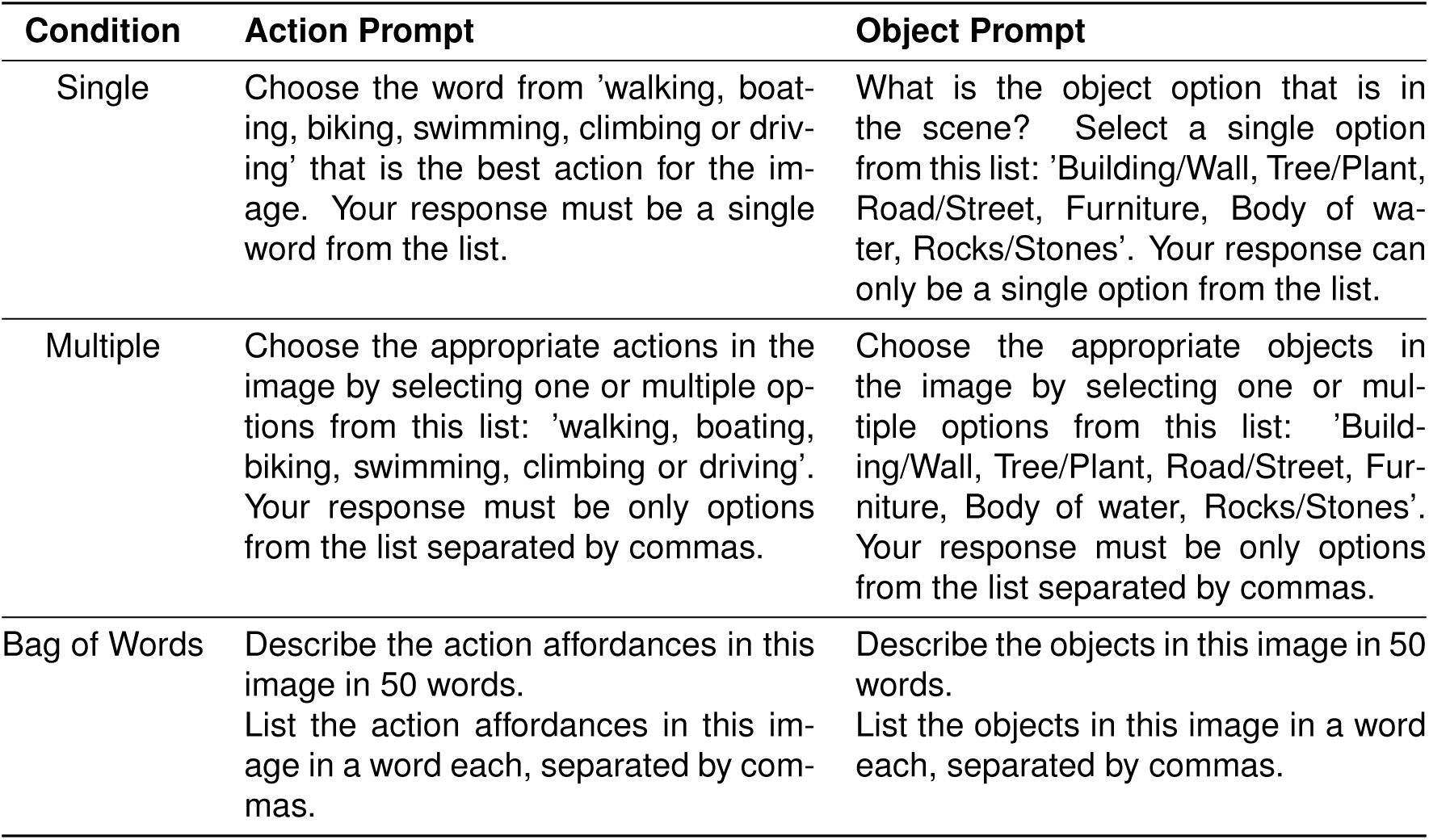
Overview of prompts used to generate GPT-4 behavioral responses for locomotive action affordances and objects. Three tasks were used: single affordance/object selection, multiple affordance/object selections, and an unrestricted (50 words) bag of words (BoW).

| Condition | Action Prompt | Object Prompt |
| --- | --- | --- |
| Single | Choose the word from 'walking, boating, biking, swimming, climbing or driving' that is the best action for the image. Your response must be a single word from the list. | What is the object option that is in the scene? Select a single option from this list: 'Building/Wall, Tree/Plant, Road/Street, Furniture, Body of water, Rocks/Stones'. Your response can only be a single option from the list. |
| Multiple | Choose the appropriate actions in the image by selecting one or multiple options from this list: 'walking, boating, biking, swimming, climbing or driving'. Your response must be only options from the list separated by commas. | Choose the appropriate objects in the image by selecting one or multiple options from this list: 'Building/Wall, Tree/Plant, Road/Street, Furniture, Body of water, Rocks/Stones'. Your response must be only options from the list separated by commas. |
| Bag of Words | Describe the action affordances in this image in 50 words.<br>List the action affordances in this image in a word each, separated by commas. | Describe the objects in this image in 50 words.<br>List the objects in this image in a word each, separated by commas. |

**Supplementary Table 4:** List of the 24 of 37 functions (affordances) attribute labels from SUN Attribute Database (Patterson et al., 2014). ** marked functions were excluded as they contained less than 300 images per category.

| Functions/affordances |  |  |
| --- | --- | --- |
| Sailing/boating | Driving | Transporting things or people |
| Biking | Vacationing/touring | Climbing |
| Sunbathing** | Reading | Teaching/training** |
| Hiking | Diving | Bathing** |
| Camping | Cleaning** | Congregating |
| Studying/learning** | Competing | Exercise |
| Research** | Gaming** | Farming |
| Swimming | Shopping | Working |
| Eating | Digging** | Praying** |
| Socializing | Using tools | Conducting business** |
| Waiting in line/queuing | Sports | Playing |
| Spectating/being in an audience | Constructing/building** | Medical activity** |

**Supplementary Table 5:** Selection of commonly used DNNs reporting the number of extracted layers and the highest correlating layer with in-scanner behavior and scene-selective ROIs. See Supplementary Table 1 for an overview of all extracted layers in each model.

| Model | Dataset | Extracted layers (Number) | Loc. Affordances | Objects | PPA | OPA | MPA |
| --- | --- | --- | --- | --- | --- | --- | --- |
| AlexNet | ImageNet | 6 | features6 | features10 | features6 | features6 | features6 |
| VGG16 | ImageNet | 14 | features0 | features26 | features0 | features0 | features0 |
| ResNet50 | ImageNet | 5 | layer2 | layer3 | layer2 | layer4 | layer2 |
| ResNet50 | Places365 | 6 | model9 | model7 | model9 | model9 | model7 |
| ResNet50 | Ade20K | 6 | decoder | decoder | decoder | decoder | decoder |
|  |  |  | ppm last conv2 | conv last1 | ppm last conv2 | ppm last conv2 | ppm last conv2 |
| DINO ResNet50 | ImageNet | 5 | layer1 | fc | layer1 | layer1 | layer1 |
| CLIP ResNet101 | ImageNet | 5 | layer3 | layer4 | layer3 | layer3 | layer3 |
| X3D-m | Kinetics | 11 | blocks.4 | blocks.4 | blocks.4 | blocks.3 | blocks.4 |
| Slowfast ResNet101 | Kinetics | 13 | blocks.5 | blocks.6 | blocks.5 | blocks.3.fast | blocks.5 |
| DINO ViT Base Patch 16 | ImageNet | 12 | blocks7attn | blocks8attn | blocks9 | blocks9 | blocks9 |
| ViT Base Patch 16 | ImageNet | 13 | blocks.11 | blocks.11 | blocks.11 | blocks.8.attn | blocks.7.attn |
| CLIP ViT-B 16 | WiT | 13 | resblocks9 | resblocks10 | resblocks7 | resblocks9 | resblocks9 |
| CLIP ViT-B 32 | WiT | 13 | resblocks10 | resblocks10 | resblocks10 | resblocks8 | resblocks8 |
| ResNet50 (finetuned) | Places365 | 52 | layer52 | layer51 | layer46 | layer52 | layer14 |
| ResNet50 (scratch) | Places365 | 51 | layer21 | layer51 | layer21 | layer21 | layer24 |
| ResNet50 (untrained) | Places365 | 52 | layer51 | layer12 | layer51 | layer51 | layer51 |

**Supplementary Table 6:** Overview of models used for fine-tuning and end-to-end trained of DNNs using affordance labels. The fine-tuned ResNet50 was pretrained on Places365, while the fine-tuned ViT Base Patch 16 was pretrained on ImageNet.

| Model | Goal | Learning Rate | Number of Epochs (5 folds) | Precision | Recall |
| --- | --- | --- | --- | --- | --- |
| ResNet50 | End-to-end training | 0.0015 | 36, 42, 42, 44, 45 | 0.73 (0.03) | 0.20 (0.04) |
| ResNet50 | Fine-tuning | 0.0053 | 33, 28, 31, 31, 34 | 0.85 (0.07) | 0.63 (0.03) |
| ViT Base Patch 16 | End-to-end training | 0.0006 | 25, 31, 28, 32, 30 | 0.68 (0.04) | 0.12 (0.03) |
| ViT Base Patch 16 | Fine-tuning | 0.0005 | 14, 19, 20, 20, 18 | 0.80 (0.01) | 0.67 (0.01) |

**Supplementary Table 7:**
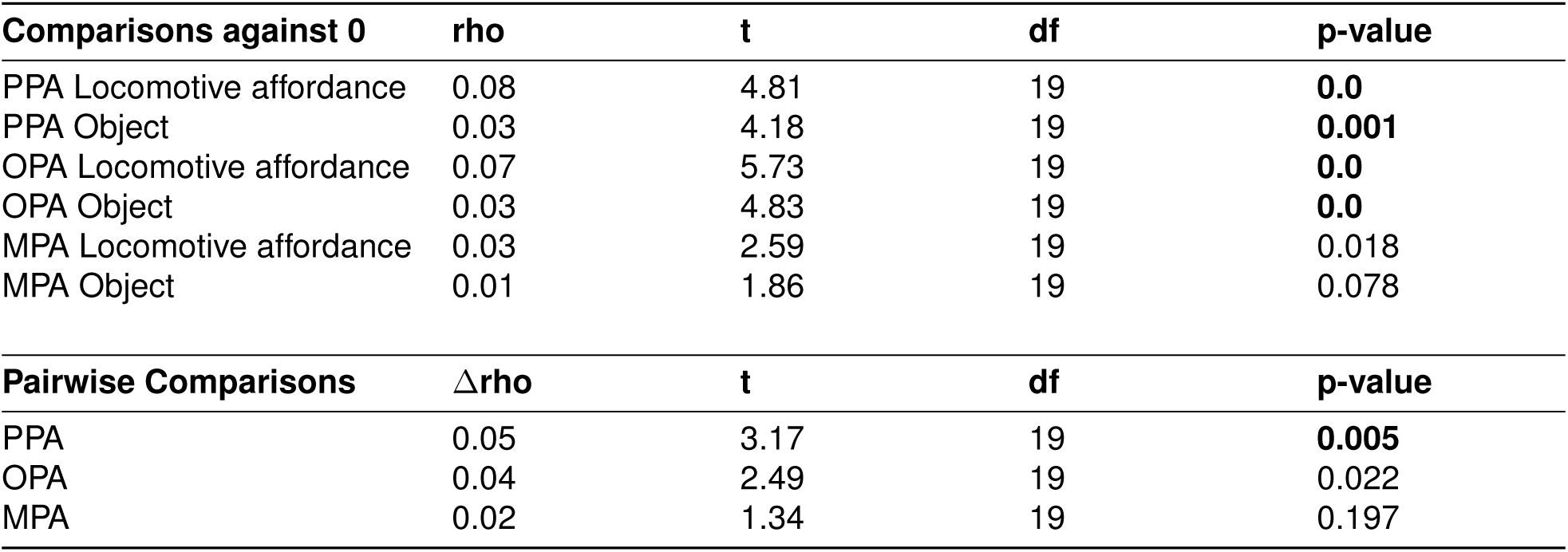
Summary of statistical results of one-sample t-test and pairwise comparisons conducted on the full correlations between action affordance annotations, object annotations versus representational similarity of fMRI responses in PPA, OPA and MPA. Rho = the average RDM correlation across subjects in each ROI (or the difference in correlation for the pairwise comparisons), t = the t-value for the one-sample t-test against zero, df = degrees of freedom and p-value = significance value. Significant p-values are highlighted in bold font only when surviving stringent Bonferroni-correction for all comparisons made within this table (here, 9 comparisons).

| Comparisons against 0 | rho | t | df | p-value |
| --- | --- | --- | --- | --- |
| PPA Locomotive affordance | 0.08 | 4.81 | 19 | **0.0** |
| PPA Object | 0.03 | 4.18 | 19 | **0.001** |
| OPA Locomotive affordance | 0.07 | 5.73 | 19 | **0.0** |
| OPA Object | 0.03 | 4.83 | 19 | **0.0** |
| MPA Locomotive affordance | 0.03 | 2.59 | 19 | 0.018 |
| MPA Object | 0.01 | 1.86 | 19 | 0.078 |
| Pairwise Comparisons | Δrho | t | df | p-value |
| PPA | 0.05 | 3.17 | 19 | **0.005** |
| OPA | 0.04 | 2.49 | 19 | 0.022 |
| MPA | 0.02 | 1.34 | 19 | 0.197 |

**Supplementary Table 8:**
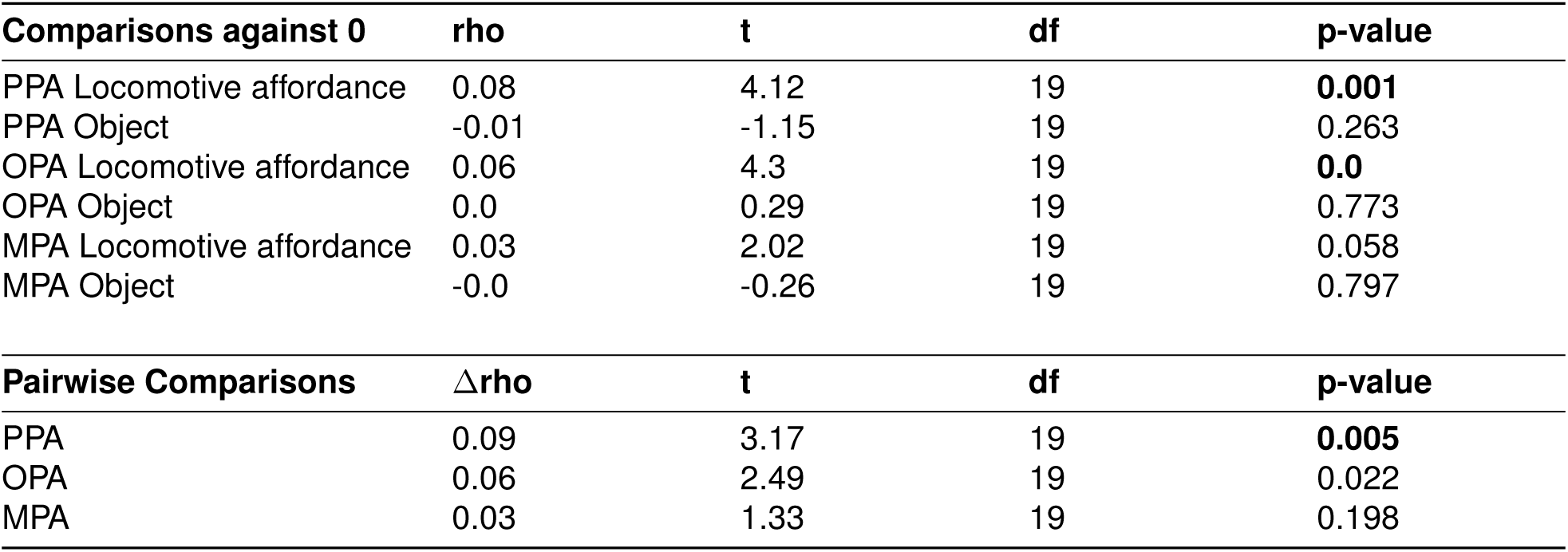
Summary of one-sample t-test and pairwise comparisons on the partial correlations between action affordance annotation, object annotations versus representational similarity of fMRI responses in PPA, OPA and MPA. See Supplementary Table 7 for details. Bold p-values indicate Bonferroni-corrected significance for all comparisons made within this table (here, 9 comparisons).

**Supplementary Table 9:**
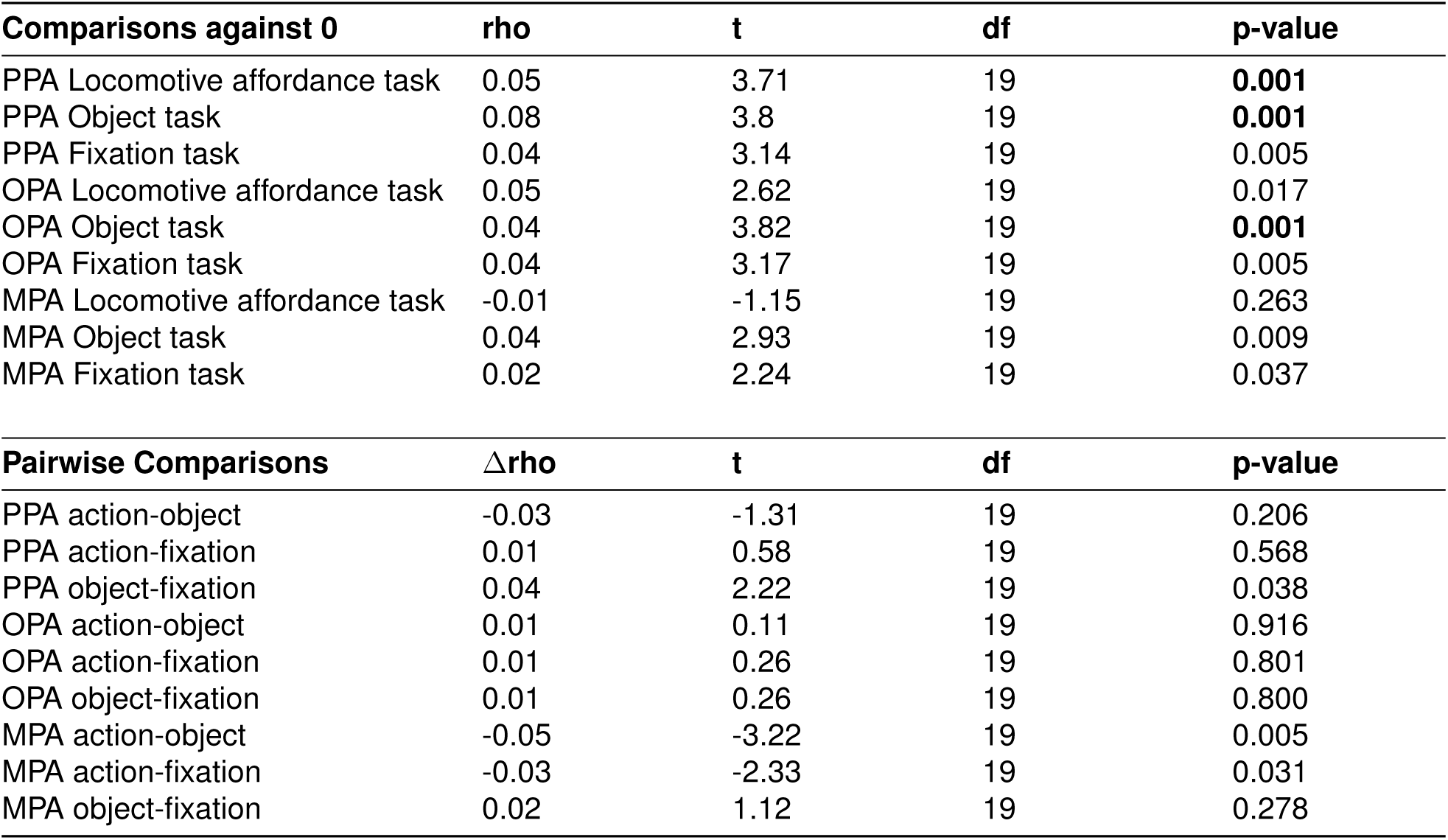
Summary of one-sample t-test and pairwise comparisons for task effect with the affordance space. Bold p-values indicate Bonferroni-corrected significance for all comparisons made within this table (here, 18 comparisons).

**Supplementary Table 10:** Summary of one-sample t-test and pairwise comparisons for task effect with the object space. Bold p-values indicate Bonferroni-corrected significance for all comparisons made within this table (here, 18 comparisons).

| **Comparisons against 0** | **rho** | **t** | **df** | **p-value** |
| --- | --- | --- | --- | --- |
| PPA Locomotive affordance task | 0.01 | 1.19 | 19 | 0.249 |
| PPA Object task | 0.03 | 4.8 | 19 | **0.0** |
| PPA Fixation task | 0.01 | 1.87 | 19 | 0.077 |
| OPA Locomotive affordance task | 0.02 | 2.77 | 19 | 0.012 |
| OPA Object task | 0.02 | 2.76 | 19 | 0.012 |
| OPA Fixation task | 0.02 | 2.82 | 19 | 0.011 |
| MPA Locomotive affordance task | 0.0 | 0.2 | 19 | 0.843 |
| MPA Object task | 0.02 | 3.13 | 19 | 0.006 |
| MPA Fixation task | 0.0 | 0.66 | 19 | 0.518 |

| **Pairwise Comparisons** | **$\Delta\rho$** | **t** | **df** | **p-value** |
| --- | --- | --- | --- | --- |
| PPA action-object | -0.02 | -2.36 | 19 | 0.029 |
| PPA action-fixation | -0.01 | -0.70 | 19 | 0.494 |
| PPA object-fixation | 0.02 | 1.52 | 19 | 0.145 |
| OPA action-object | -0.01 | -0.12 | 19 | 0.905 |
| OPA action-fixation | -0.01 | -0.73 | 19 | 0.475 |
| OPA object-fixation | -0.01 | -0.68 | 19 | 0.506 |
| MPA action-object | -0.02 | -1.86 | 19 | 0.078 |
| MPA action-fixation | -0.01 | -0.20 | 19 | 0.844 |
| MPA object-fixation | 0.02 | 1.71 | 19 | 0.104 |

**Supplementary Table 11:** Summary of one-sample t-test and pairwise comparisons for the partial correlations between CLIP ViT-B 32 and the locomotive affordances and object space in the ROIs. Bold p-values indicate Bonferroni-corrected significance for all comparisons made within this table (here, 18 comparisons).

| **Comparisons against 0** | **rho** | **t** | **df** | **p-value** |
| --- | --- | --- | --- | --- |
| PPA Locomotive affordance | 0.07 | 3.87 | 19 | **0.001** |
| PPA CLIP ViT-B 32 (WiT) | 0.03 | 5.11 | 19 | **0.0** |
| OPA Locomotive affordance | 0.06 | 4.17 | 19 | **0.001** |
| OPA CLIP ViT-B 32 (WiT) | 0.03 | 4.9 | 19 | **0.0** |
| MPA Locomotive affordance | 0.02 | 1.61 | 19 | 0.125 |
| MPA CLIP ViT-B 32 (WiT) | 0.02 | 4.45 | 19 | **0.0** |
| **Pairwise Comparisons** | **Δrho** | **t** | **df** | **p-value** |
| PPA | 0.04 | 1.76 | 19 | 0.094 |
| OPA | 0.03 | 1.31 | 19 | 0.205 |
| MPA | 0.01 | -0.19 | 19 | 0.854 |
| **Comparisons against 0** | **rho** | **t** | **df** | **p-value** |
| PPA Object | -0.01 | -1.61 | 19 | 0.124 |
| PPA CLIP ViT-B 32 (WiT) | 0.05 | 7.53 | 19 | **0.0** |
| OPA Object | 0.0 | 0.2 | 19 | 0.844 |
| OPA CLIP ViT-B 32 (WiT) | 0.04 | 7.55 | 19 | **0.0** |
| MPA Object | -0.01 | -1.37 | 19 | 0.186 |
| MPA CLIP ViT-B 32 (WiT) | 0.03 | 5.44 | 19 | **0.0** |
| **Pairwise Comparisons** | **Δrho** | **t** | **df** | **p-value** |
| PPA | -0.06 | -4.92 | 19 | **0.0** |
| OPA | -0.04 | -3.29 | 19 | 0.004 |
| MPA | -0.04 | -3.22 | 19 | 0.004 |

